# Single-cell spatial analysis of pediatric high-grade glioma reveals a novel population of SPP1^+^/GPNMB^+^ myeloid cells with immunosuppressive and tumor-promoting capabilities

**DOI:** 10.1101/2025.03.18.643953

**Authors:** Thijs J.M. van den Broek, Raoull Hoogendijk, Mariëtte E.G. Kranendonk, Julie A.S. Lammers, Akshaya L. Krishnamoorthy, Ravian L. van Ineveld, Milo Molleson, Vasily O. Tsvetkov, Femke C. A. Ringnalda, Marc van de Wetering, Yan Su, John I. Bianco, Cristian Ruiz Moreno, Mario G. Ries, Eelco W. Hoving, Jasper van der Lugt, Leila Akkari, David P. Schrijver, Hendrik G. Stunnenberg, Anne C. Rios, Dannis G. van Vuurden, Anoek Zomer

## Abstract

**Background:** Pediatric-type diffuse high-grade gliomas (pHGG) are a leading cause of pediatric cancer-related mortality. Although immunotherapy offers a promising treatment avenue, clinical responses in pHGG patients remain limited. A detailed understanding of the tumor immune microenvironment (TIME) is essential for advancing immunotherapeutic strategies.

**Methods:** We performed single-cell spatial analysis integrating cyclical immunofluorescence imaging and Spatial Molecular Imaging to interrogate the proteomic and transcriptomic landscape of pHGG. A tissue microarray comprising 32 diagnostic patient-derived pHGG samples was utilized to map the spatial distribution of immune and tumor cells.

**Results:** Our analyses reveal that the pHGG TIME is predominantly composed of myeloid cells, including brain-resident microglia and monocyte-derived macrophages, with only few T cells. A significant subset of these myeloid cells express mesenchymal-like genes and are positive for SPP1 and GPNMB. Spatial mapping further demonstrated that SPP1^+^/GPNMB^+^ myeloid cells localize in close proximity to mesenchymal-like tumor cells, and negatively correlate with the location and presence of CD8^+^ T cells. These cells also express genes related to immunosuppression and epithelial-to-mesenchymal transition, indicating their potential role in establishing an immunosuppressive niche.

**Conclusions:** Our findings reveal a distinct immune landscape in pHGG characterized by SPP1^+^/GPNMB^+^ myeloid cells which may contribute to the exclusion of CD8^+^ T cells. This spatially resolved insight identifies these myeloid cells as promising therapeutic targets and provides a rationale for developing novel immunotherapeutic strategies to improve outcomes in pediatric high-grade gliomas.

## Introduction

Pediatric-type diffuse high-grade gliomas (pHGG) are among the most aggressive and lethal types of tumors, with a median overall survival of just 18 months for hemispheric tumors and 13.5 months for midline tumors^1^. Recent advances in immunotherapies have shown promising results for several brain tumors, including pHGG^2,3,4^. However, response rates vary largely between patients. It is hypothesized that the efficacy of these therapies is negatively influenced by an immunosuppressive tumor-immune microenvironment (TIME), underscoring that a comprehensive understanding of the TIME in pHGG is required. Studies have shown that a significant fraction of the pHGG TIME consists of macrophages, including brain-resident microglia (MG) and monocyte-derived macrophages (MDMs), with only low levels of tumor-infiltrating lymphocytes^5,6,7,8,9^. Macrophages are highly plastic and can adopt different phenotypes depending on environmental conditions^10,11^. Although *in vitro* experiments led to a dualistic paradigm describing two opposing phenotypes, immune-stimulatory M1 and immune-inhibitory M2 macrophages, a continuum of macrophage states is found *in vivo* where cells are exposed to a multitude of signals from the environment^12^. Immune-inhibitory myeloid cells can contribute to an immunosuppressive TIME which prevents the influx and/or activation of lymphocytes such as cytotoxic T cells, observed in other cancers^13,14^. However, whether and how myeloid cells contribute to low levels of lymphoid influx in pHGG remains unclear. The field lacks knowledge of the phenotypic diversity of macrophages in pHGG and the potential functional consequences of this heterogeneity. Additionally, cellular interactions and their spatial context within the TIME have not been characterized for pHGGs. The spatial organization of the TIME has been linked to tumor growth and progression in several studies, highlighting the clinical value of spatial analyses^15,16^. A thorough understanding of cellular interactions and the spatial organization of the TIME might predict response to cellular immunotherapies, as has been shown for other tumor types^17,18^.

Here, we used cyclical immunofluorescence imaging (cIF) in combination with Spatial Molecular Imaging (SMI) to study the pHGG TIME and its spatial organization at a proteomic and transcriptomic level. We observed differences in the composition of the TIME between pHGGs found in the midline compared to non-midline pHGGs; however, in all pHGGs, the TIME was dominated by myeloid cells, with low amounts of T cells. A significant percentage of myeloid cells exhibited a particular phenotype characterized by high *SPP1* and *GPNMB* expression. SPP1^+^/GPNMB^+^ myeloid cells were further characterized by an accumulation of lipid droplet proteins and increased expression of lipid metabolism-related genes. Our findings suggest a reciprocal interaction between SPP1^+^/GPNMB^+^ myeloid cells and MES-like tumor cells. This may be important clinically as we observed these myeloid cells to be associated with immunosuppression, exemplified by reduced CD8^+^ T cell proximity and local upregulation of immunosuppressive cytokine and chemokine signaling. Together, these results reveal spatial immune-tumor heterogeneity and local immunosuppression with a major role for SPP1^+^/GPNMB^+^ myeloid cells, offering new leads to improve immunotherapy targeting pHGG.

## Results

### Mapping the tumor microenvironment of pHGG using cyclical immunofluorescence imaging

To spatially map the immune compartment of pHGG, we generated a TMA containing 32 formalin-fixed paraffin-embedded (FFPE) pHGG tumor samples collected at diagnosis (Fig. 1A, Supplementary Table 1). Among these were 14 diffuse midline gliomas H3 K27-altered (DMG), 11 diffuse pediatric-type high-grade glioma H3-wildtype and IDH-wildtype, 3 infant-type hemispheric gliomas, 3 diffuse hemispheric gliomas H3 G34-mutant, and 1 diffuse hemispheric glioma, H3 K27-altered not elsewhere classified. We applied cIF on the TMA using a panel specifically designed to identify major immune cell types, endothelial cells, and myeloid cell phenotypes. We observed substantial differences in immune cell frequencies (Fig. 1C), particularly when comparing DMGs to non-midline-located pHGGs (NM-HGGs). With one exception, myeloid cells were by far the largest immune component in all tumors analyzed. Interestingly, we found that the percentages of brain-resident MG and peripherally recruited MDMs varied according to the patient group, with more MDMs in NM-HGGs (Fig. 1D). Furthermore, we observed a significantly higher percentage of T cells in NM-HGGs compared to DMG (1.57% vs. 0.48%, Fig. 1E), in line with previous publications^5,7^. The increased levels of peripheral immune cells in NM-HGGs could not be explained by the level of vascularization, which was found to be similar in DMG and NM-HGG (Fig. 1F).

**Figure 1.**
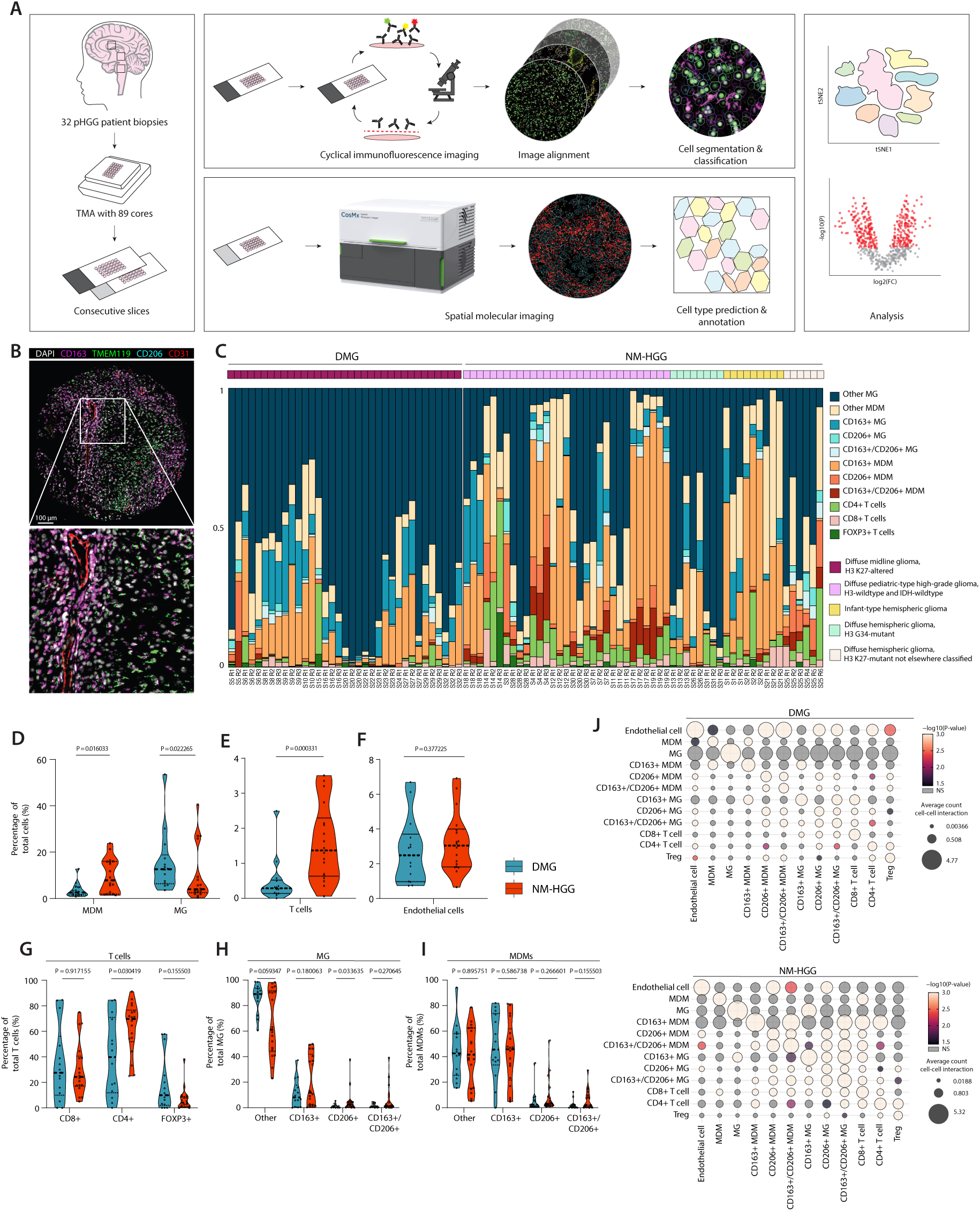
Mapping of pHGG patient biopsies using cyclical immunofluorescence and spatial molecular imaging. A, Schematic overview of the current study. **B,** Representative immunofluorescence image of a TMA core from a panel targeting 10 proteins. DAPI, CD163, TMEM119, CD206, and CD31 are visualized in this image. **C,** Stacked bar plot showing the fraction of immune cells detected per TMA core. Each bar represents one TMA core. Tumor classification according to the 2021 WHO classification of CNS tumors^22^ is shown above the stacked bar plot. **D, E, F,** Violin plots comparing the frequencies of immune cell types as a percentage of total cells detected per patient between DMG (n=15) and NM-HGG (n=17). Each dot represents one patient. P-values determined by Mann-Whitney U tests. **G, H, I,** Violin plots comparing the frequency of immune cell subsets within T cells (**G**), MG (**H**), and MDMs (**I**), between DMG (n=15) and NM-HGG (n=17). P-values determined by Mann-Whitney U tests. **J,** Dot plot visualizing the spatial cellular co-localization (and interaction) as determined by neighborhood permutation testing within the imcRtools package^23^ . A significance threshold of P = 0.05 was used.

Next, we made use of the different markers included in our cIF experiment to subset T cells. In both DMG and NM-HGG, most T cells showed expression of CD4 and were negative for CD8 and FOXP3 (Fig.1G), confirming previous observations made using classical immunohistochemistry^7^. Although more T cells were detected within NM-HGGs than in DMGs, there was no significant difference in the proportion of CD8^+^ T cells between the two groups. However, the CD4 fraction of T cells was significantly lower in DMGs while the proportion of regulatory T cells (Tregs), characterized by FOXP3 expression, was similar compared to NM- HGGs.

To determine the phenotypic state of myeloid cells in the pHGG environment, we assessed the expression of CD163 and CD206 on MG and MDMs separately (Fig. 1H, I). These scavenger receptors are considered classical markers for immunosuppressive myeloid populations^19^. MG were discriminated from MDMs based on the expression of TMEM119. In both DMG and NM-HGG, the vast majority of MG was negative for both CD163 and CD206 (Fig. 1H). We also found a large proportion of double-negative MDMs in both patient groups, with CD163^+^/CD206^-^ MDMs being the other major phenotype (Fig. 1I). CD163^+^/CD206^+^ MG and MDMs represent only a small fraction of all MG/MDMs. Interestingly, more CD163^-^/CD206^+^ cells were found in NM-HGGs compared to DMG. These myeloid cells likely represent central nervous system (CNS) border-associated myeloid cells characterized by high CD206 expression^20,21^. Indeed, a comparative analysis of distances between myeloid cells and the nearest endothelial cells revealed that both MG and MDMs expressing CD206 are closely associated with the vasculature in both DMG and NM-HGG (Supplementary Fig. 1C, D).

To further characterize patterns of communication between individual cell types and phenotypes, we quantified cell-cell colocalization within pHGG tissues through neighborhood permutation testing. Our data revealed that in DMGs, CD8^+^ T cells were predominantly found to interact with MG rather than MDMs, an observation that we confirmed through single-cell distance analysis (Fig 1J, Supplementary Fig. 1D). In contrast, in NM-HGGs, CD8^+^ T cells were found close to both MG and MDMs. Unexpectedly, despite the high abundance of CD163^-^/CD206^-^ MG and MDMs relative to the other phenotypes (Fig. 1H, I), CD8^+^ T cells were mainly found to colocalize with myeloid cells expressing both or one of the immunosuppressive markers, in both patient groups (Fig. 1J). In addition, another potential type of immunosuppressive cell-cell communication was identified in the form of Tregs, predominantly interacting with CD4^+^ T cells in both DMG and NM-HGG.

In summary, our data confirms that myeloid cells are the most abundant immune component in the TIME of both DMG and NM-HGG, and that most of these cells lack expression of the classic immunosuppressive markers CD163 and CD206. The few CD8^+^ T cells found in these tumors preferentially interact with myeloid cells positive for both or one of these suppressive markers. Specifically for DMG, we observed less systemically recruited immune cells (T cells and MDMs), and CD163^-^/CD206^+^ MG/MDMs.

### Spatial transcriptomics reveals the presence of MES-like myeloid cells in pHGG

To comprehend the spatial composition and immune-regulatory mechanisms of pHGG in more detail while maintaining single-cell resolution, we performed a Spatial Transcriptomic experiment employing SMI of 1000 RNA probes using the NanoString CosMx platform. For this, we used the TMA (Fig. 1A, Supplementary Fig. 2A), and analyzed multiple DMGs and NM-HGGs in parallel. We observed a significant correlation between protein expression in our cIF data and RNA expression in the spatial transcriptomic data (Supplementary Fig. 2A-C). We identified the previously described transcriptionally defined tumor cell types, including astrocyte (AC)-like, oligodendrocyte (OC)-like, oligodendrocyte precursor (OPC)-like, mesenchymal (MES)-like, and neural-progenitor (NPC)-like cells^24,25,26^ (Fig. 2A, B). In addition, we detected the major immune cell populations, as well as a small population of B cells in three NM-HGG samples (Fig. 2C, D).

**Figure 2.**
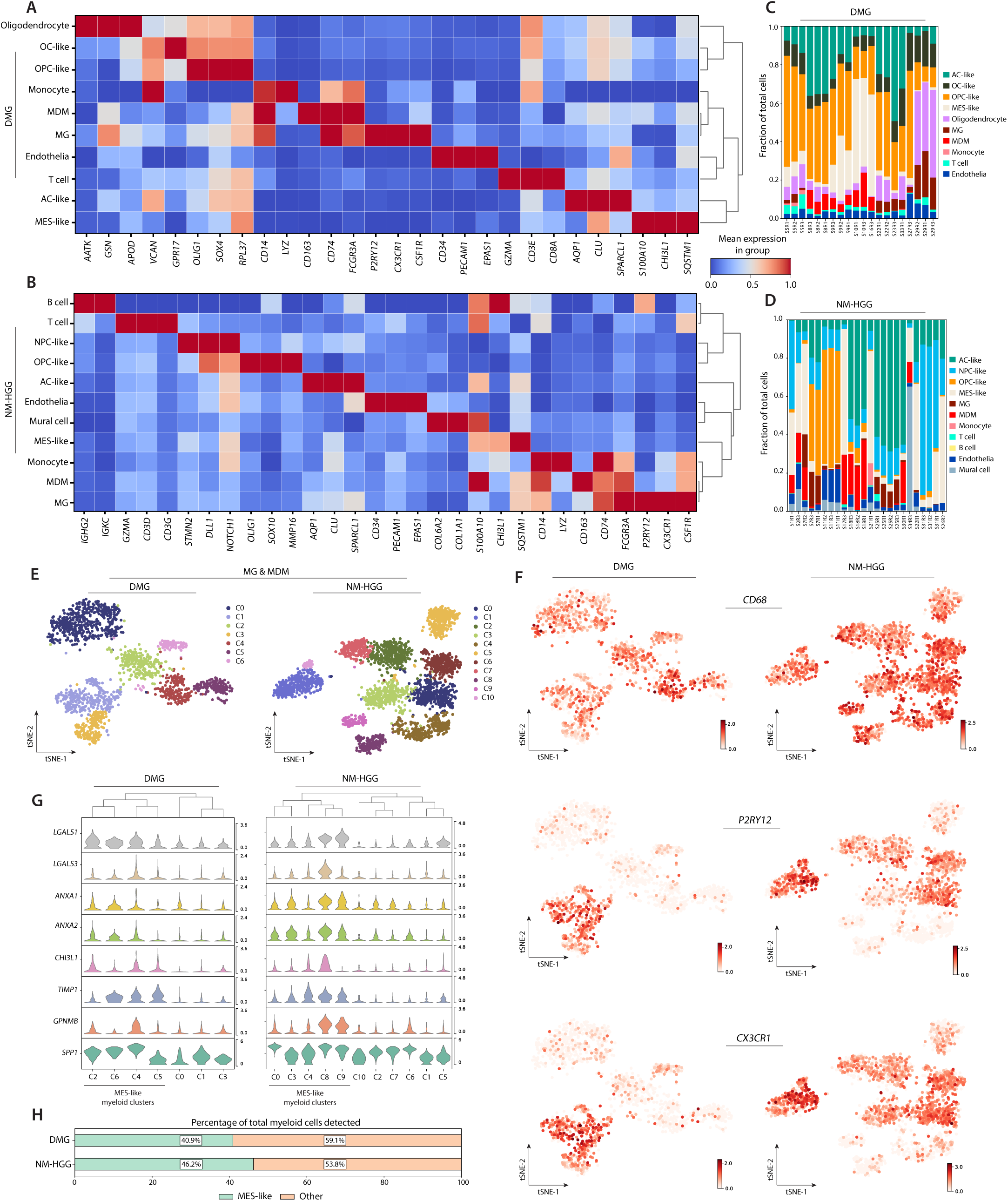
Single-cell spatial transcriptomics reveals the presence of MES-like myeloid cells in pHGG. **A**, **B**, Heatmap of the top markers for each cell type annotated within DMG (**A**) and NM-HGG (**B**) using SMI on the Nanostring CosMx. The scaled mean expression within each group is indicated by the color scale. The dendrogram represents hierarchical clustering based on Pearson correlation across principal component analysis (PCA) components. **C**, **D**, Stacked bar charts showing the cell frequencies as a fraction of total cells detected in each field of view (FOV) for DMG (**B**) and NM-HGG (**D**). Each FOV was captured at the center of a TMA core. **E**, t-distributed Stochastic Neighbor Embedding (tSNE) visualization of the MG and MDM cell populations within DMG and NM-HGG. Color coding specifies the clusters identified using the Leiden algorithm^33^. **F**, tSNE plots for the MG and MDM cell populations of DMG and NM-HGG with the expression of general myeloid gene CD68, and microglia- specific genes P2RY12 and CX3CR1 visualized by color. **G,** Violin plots visualizing the expression of MES-like genes across myeloid clusters detected. High expression of GPNMB and SPP1 were characteristic of the identified MES-like myeloid clusters. **H,** Stacked bar plot showing the MES-like myeloid cells as a percentage of total myeloid cells detected by spatial transcriptomics.

Similar to our cIF data (Fig. 1C), we observed substantial differences in both neoplastic and immune cell frequencies between samples of different patients (Fig. 2C, D). In line with our previous analysis, myeloid cells were found to be the main component of the TIME in all samples analyzed. We therefore decided to redirect our analyses towards further investigating MG and MDMs, and performed a Leiden clustering analysis where we detected 6 clusters in DMG and 10 clusters in NM-HGGs (Fig. 2E). We were able to differentiate between the MG and MDM clusters based on expression of *P2RY12* and *CX3CR1,* classical MG marker genes (Fig. 2F).

In several myeloid clusters, we noticed similar expression patterns between DMGs and NM-HGGs (C2, C4, C5, C6 in DMGs, and C0, C3, C4, C8, C9 in NM-HGGs). These clusters showed high expression of genes such as *CHI3L1*, *ANXA1*, and *LGALS1* (Fig. 2G), typically reported in the context of epithelial-mesenchymal transition (EMT) and found to be expressed in MES-like tumor cells^26^. Therefore, we termed these myeloid cells MES-like myeloid cells. The identified MES-like myeloid clusters were characterized by high expression of *SPP1* and *GPNMB* (Fig. 2G). Differential gene expression (DGE) analysis showed that the expression of these genes was significantly higher in MES-like myeloid clusters than in other myeloid clusters in both DMGs and NM-HGGs (Supplementary Fig. 2D). Consequently, we will use these markers, along with myeloid markers, to identify MES-like myeloid cells in subsequent analyses.

Since MES-like myeloid cells comprised a significant part of the total myeloid cells detected (40.9% for DMG and 46.2% for NM-HGG; Fig. 2H), we hypothesize that they may have a major role in the pHGG TIME. Indeed, recent evidence suggests a pivotal role for secreted phosphoprotein 1 (SPP1) and/or glycoprotein non-metastatic B (GPNMB) under pathological conditions. SPP1 has been described in a broad range of disease-associated myeloid cells, such as damage-associated MG during brain injury^27^. Recent studies have also pointed towards an immunosuppressive role of SPP1 in myeloid cells^28,29^. Similarly, roles in immunosuppression and EMT-induction have been outlined for GPNMB-expressing macrophages in murine models of adult glioblastoma^30,31^. Additionally, Kloosterman *et al*. recently revealed that GPNMB^+^ lipid-accumulating myeloid cells in adult glioblastoma fuel the proliferation of cancer cells through the transfer of lipids^32^.

### Pro-tumorigenic interaction networks between myeloid cells and MES-like tumor cells

To further characterize the MES-like myeloid clusters, we performed single-cell distance analysis and revealed MES-like myeloid cells to spatially co-occur with MES-like tumor cells in both DMG and NM-HGGs (Fig. 3A, B). To validate this finding, we collected bulk RNA-sequencing data from an extended cohort of patients diagnosed with pHGG, including 31 DMGs and 39 NM-HGGs. Within this dataset, we generated MES-like signature scores and OPC/OC-like signature scores for each sample. We observed a significant positive correlation between *GPNMB* expression, and a MES-like signature score for both DMGs and NM-HGGs (Fig. 3C). *GPNMB* expression negatively correlates with OPC/OC-like signatures (Supplementary Fig. 3A, B). Next, we split the DMGs and NM-HGGs into a *GPNMB*-low group and a *GPNMB*-high group and performed DGE analysis. We observed a statistically significant upregulation of the MES-like genes *CHI3L1*, *LGALS1*, and *ANXA2* in the *GPNMB-*high group for both DMGs and NM-HGGs (Fig. 3D). Other upregulated genes include *MARCO* and *S100A4*, two markers that have been reported as potential targets for immunotherapy in adult glioma^34,35^. The scavenger receptor *MARCO* is commonly found on lipid-laden macrophages^36^. Furthermore, we observed *CXCL8* (linked to immunosuppressive myeloid cell recruitment in other cancers^37^), the IL1 decoy receptor *IL1R2,* and *LIF* (associated with negative regulation of CD8^+^ T cell infiltration^38^) to be upregulated in the *GPNMB*-high group. The matrix metalloproteinases *MMP10* and *MMP12* have both been described in the context of cancer cell invasion and EMT^39,40^. Finally, we observed upregulation *PLIN2* which translates for perilipin-2, a protein that binds to lipid droplets as they accumulate within cells^41^.

**Figure 3.**
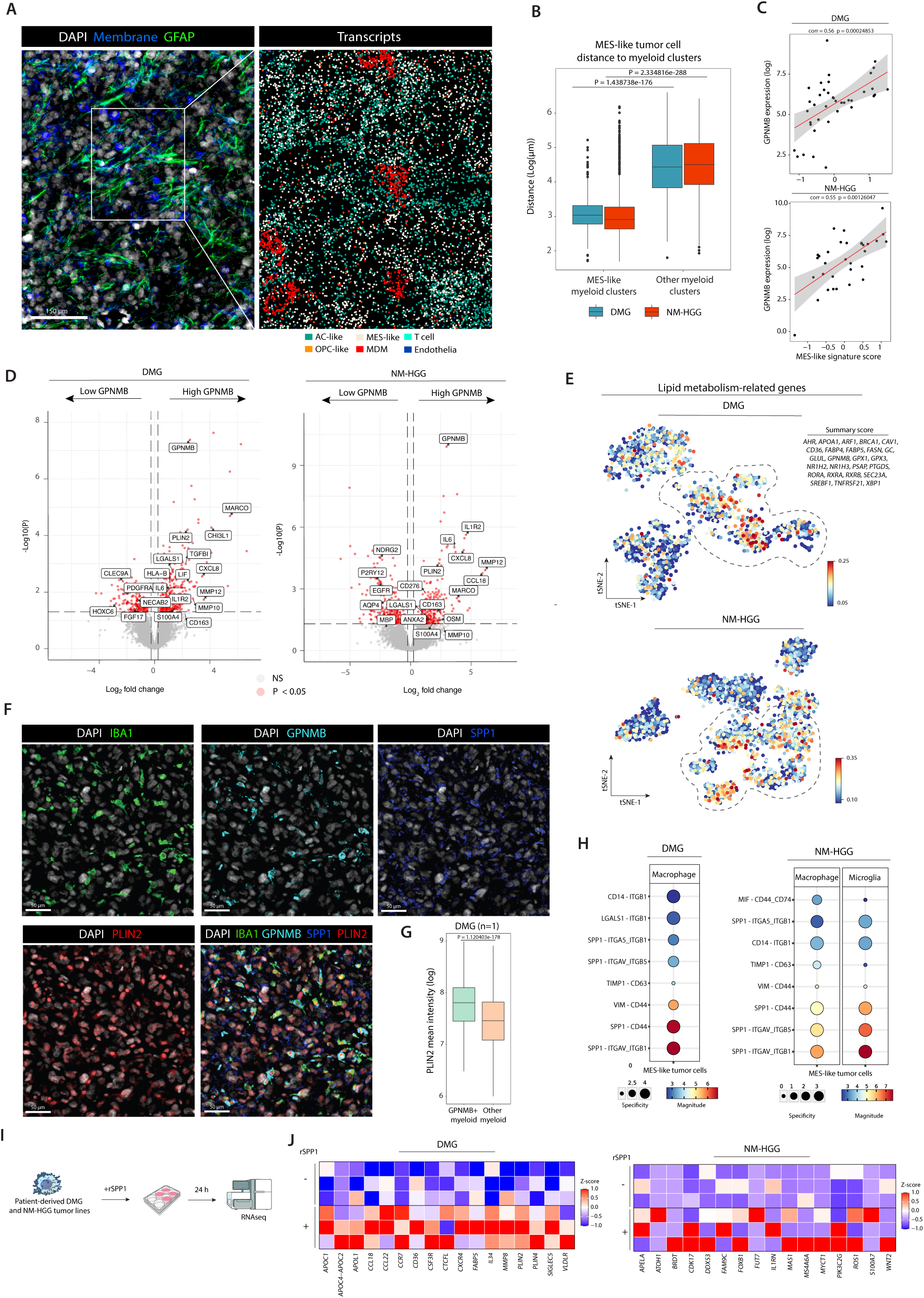
MES-like myeloid cells display increased lipid metabolism and lipid accumulation. **A**, Representative immunofluorescence image (top; membranes are visualized by CD298/B2M/18s rRNA) and detected transcripts capture by the Nanostring CosMx (bottom) within a MES-like tumor region. Transcripts in the bottom image are color-coded by cell type. **B,** Box plot comparing the distance of MES-like tumor cells to the nearest MES-like myeloid cells or other myeloid cells. P-values determined by Mann-Whitney U tests. Dots represent outliers. **C,** Scatter plots showing the correlation between GPNMB expression and a MES-like signature score using bulk RNA-seq data from NM-HGG patients (n=31) and DMG patients (n=39). Each dot represents on patient. P-values and correlation determined by Spearman correlation testing. **D,** Volcano plots of differentially expressed genes between patients with low GPNMB expression and patients with high GPNMB expression. **E,** tSNE plots for the MG and MDM cell populations of DMG and NM-HGG color-coded for a summary score calculated for lipid metabolism-related genes. MES-like myeloid clusters are indicated with a grey dashed line. **F,** Representative immunofluorescence image of a DMG sample stained for IBA1, GPNMB, SPP1, and PLIN2. **G,** Box plot comparing the mean intensity of PLIN2 detected within GPNMB^+^ myeloid cells compared to other (GPNMB^-^) myeloid cells within the DMG sample of (**E**). P-value determined by Mann-Whitney U test. **H,** Dot plots visualizing ligand-receptor interactions within FOVs captured by the CosMx which were enriched for MES-like tumor cells. For DMGs, only macrophages were detected in MES-like tumor- enriched regions, for NM-HGGs both macrophages and microglia. **I,** Three DMG (HSJD-DIPG-007, HDJS-DIPG-012, M354AAB- TO) and three NM-HGG (HSJD-GBM-001, HSJD-GBM-002, KNS-42) patient-derived tumor lines were profiled by bulk RNA sequencing after 24 h of exposure to recombinant SPP1 protein. **J,** Heatmaps showing Z-scored expression of DESEQ2 normalized counts of genes of interest after the experiment shown in (**I**).

To investigate whether the MES-like myeloid cells detected in pHGG patients are similar to the lipid-accumulating myeloid cells found in adult glioma^32^, we determined the expression score of lipid metabolism-related genes (such as *FABP4*, *FABP5*, *CD36*, *PSAP, and PTGDS*) within all myeloid clusters in the CosMx data. Indeed, we found elevated levels of these genes in the MES-like myeloid clusters expressing *GPNMB* (Fig. 3E). The combined expression of MES-like genes and lipid-metabolism genes which we detected in pHGG myeloid cells aligns with myeloid signatures described in adult glioma^32^, and a wide range of other pathologies, including Alzheimer’s disease^42^, and multiple sclerosis^43^. To determine whether the upregulation of these genes in MES-like myeloid cells is functional, we assessed the degree of lipid accumulation within these cells by staining for PLIN2. We observed PLIN2 expression to be significantly higher in GPNMB^+^ myeloid cells compared to other myeloid cells within a full DMG tissue section (Fig. 3F and G). This indicates that these myeloid cells indeed exhibit increased uptake of lipids, similar to lipid-laden macrophages in glioblastoma^32^. The accumulation of lipid droplets within CNS myeloid cells and the upregulation of PLIN2 have been described in demyelinating disorders such as multiple sclerosis^44^. Furthermore, SPP1^+^ microglia have been described as an important factor of axonal demyelination in the context of pontine infarction, which might contribute to the widespread pontine axonal degeneration and resulting clinical deterioration which is often observed in pontine DMG patients at end- stage disease^45^.

To determine how the MES-like myeloid cells interact and possibly influence MES-like tumor cells, we performed LR interaction analysis between MDM/MG and MES-like tumor cells in the FOVs of the CosMx data enriched for MES-like tumor cells. This showed significant interactions between *SPP1* on myeloid cells and its receptor *CD44* on tumor cells in both DMGs and NM-HGGs (Fig. 3H). *CD44* is a marker found to be upregulated in glioma cells with a MES-like state ^26^, and signaling through this receptor is known to promote their invasive and proliferative capacity^46,47^. Integrins expressed by the MES-like tumor cells were identified as another interaction partner for proteins expressed on myeloid cells, including *SPP1* and *LGALS1*. Integrin signaling in cancer is widely known to be pro-mitogenic and pro-migratory^48^. Additionally, myeloid cell-expressed *TIMP1* showed a significant interaction with *CD63* expressed on tumor cells in all pHGGs. *TIMP1-CD63* interactions have been observed to induce EMT via mesenchymal protein markers^49^.

To further explore how MES-like myeloid cells may influence the phenotypic state of tumor cells, we cultured three DMG (H3 K27-altered) and three NM-HGG (H3G34-mutant, and H3, IDH-wildtype) patient-derived tumor lines in the presence of recombinant SPP1 (Fig. 3H). After 24 hours of exposure, we performed bulk RNA-sequencing and compared profiles of SPP1-treated cells with untreated controls (Fig. 3I). Interestingly, we observed an increased expression of lipid metabolism-related genes (such as *APOC1*, *APOL1*, *PLIN2*, *PLIN4*, and *VLDLR*) in the three DMG lines, but not in the NM-HGG cell lines (Fig. 3J). This observation supports the hypothesis of metabolic crosstalk facilitating lipid uptake and exchange between tumor cells and lipid-accumulating myeloid cells in adult glioma^32^. In NM-HGG, we observed increased expression of several genes linked to cell proliferation (*BRDT*, *CDK17)*^50,51^, migration (*PIK3C2G*, *APELA*)^52,53^, and stemness (*MYCT1, ROS1, WNT2)*^54,55,56^ (Fig. 3J). The distinct responses observed between DMG and NM-HGG tumor cells likely reflect the unique tumor biology of these subgroups, which is further influenced by their differing anatomical locations and genetic profiles.

Altogether, our findings suggest that SPP1^+^/GPNMB^+^ myeloid cells correlate with MES-like tumor states and that reciprocal interactions between these myeloid cells and tumor cells may induce pro-tumorigenic pathways within tumor cells.

### SPP1^+^/GPNMB^+^ regions are characterized by immunosuppressive signaling

Given that our bulk RNA-seq analysis identified several immunosuppressive genes correlating with high *GPNMB* expression (Fig. 3D) and that lipid-laden macrophages have been associated with immunosuppression in other cancers^57^, we sought to investigate the immunosuppressive characteristics of the SPP1^+^/GPNMB^+^ myeloid cells. To that end, we utilized a published *in vitro* model for lipid-laden macrophages^58^. We isolated peripheral blood-derived monocytes from healthy adult donors, which we differentiated into naïve macrophages and subsequently exposed to myelin debris (Fig. 4A). The phenotype of the macrophages was confirmed by flow cytometry using GPNMB and the lipid marker BODIPY (Fig. 4B). Increased lipid uptake was found to coincide with an increase in GPNMB expression (Fig. 4B, C). Using bulk RNA-seq, we analyzed transcriptomic changes upon exposure to myelin debris, as well as in combination with tumor-conditioned medium (TCM) collected from patient-derived tumor cells. This revealed upregulation of lipid metabolism-related genes in the myelin-exposed conditions (Fig. 4D, Supplementary Fig. 4B). Combining myelin with tumor-secreted factors induced expression of these genes even further. In addition, macrophage exposure to myelin or myelin combined with TCM led to increased expression of immunosuppressive cytokines and chemokines. Among the upregulated chemokines were several immunosuppressive genes such as *CXCL16*, *CXCL8*, and *TNFSF14*, which have been linked to immunosuppressive myeloid cell recruitment in other cancers^59,37,60^. Furthermore, *EBI3* (a known inducer of Treg development^61^), *GRN* (associated with reduced CD8^+^ T cell infiltration^62^), *CD244* (expressed on myeloid-derived suppressor cells^63^), *IL10RA* (regulates anti-inflammatory macrophage function^64^, and *LIF* (also identified in Figure 3D) were found to be upregulated after exposure to myelin and TCM.

**Figure 4.**
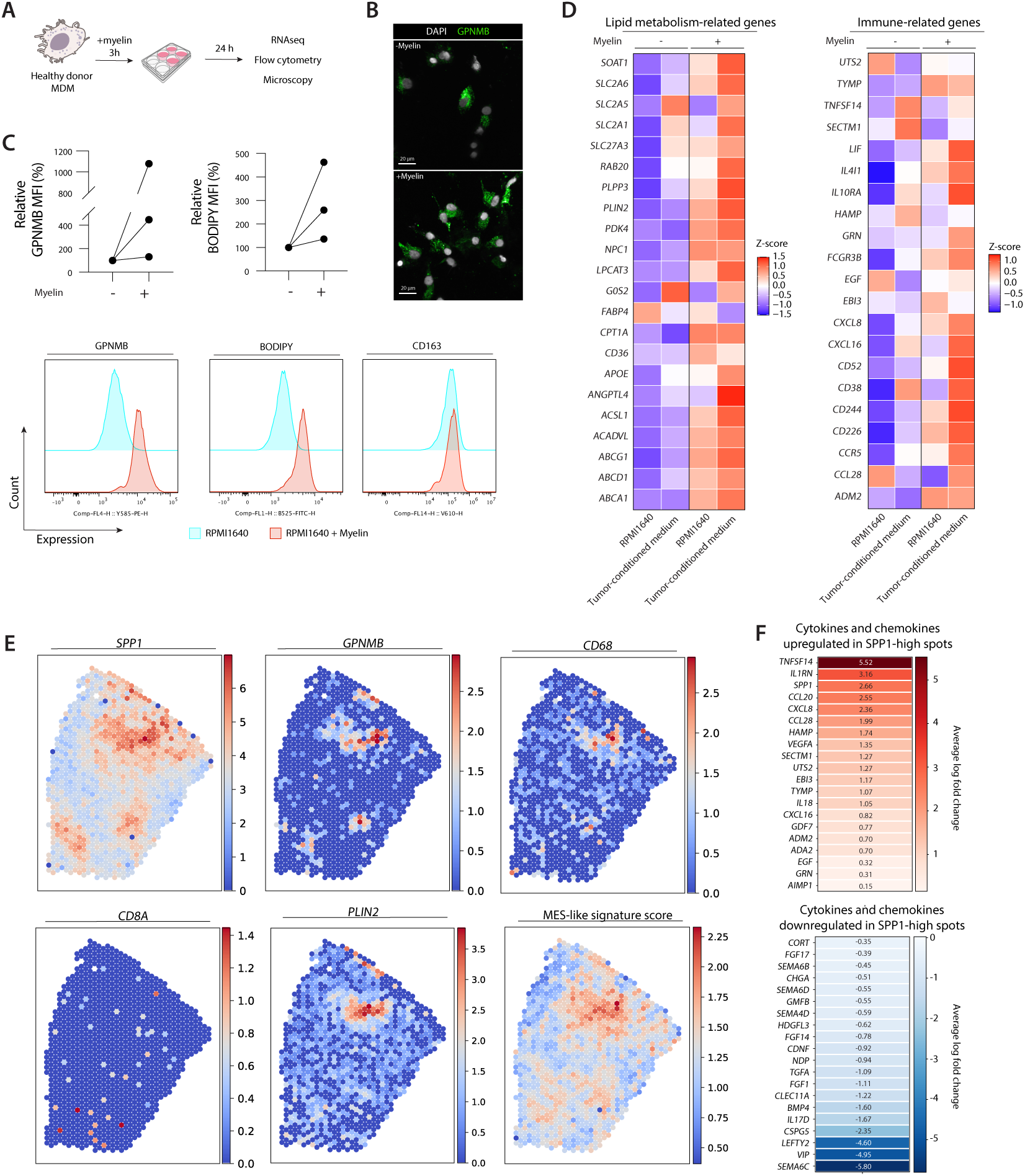
SPP1+/GPNMB+ regions are characterized by immunosuppressive signaling. A, In vitro setup to model SPP1^+^/GPNMB^+^ myeloid cells. **B,** Representative immunofluorescence image of in vitro cultured macrophages showing protein expression of GPNMB. **C,** Flow cytometry analysis of macrophages exposed to mouse-derived myelin to determine expression of GPNMB, CD163, and BODIPY (n=3 donors). **D,** Macrophages (n=3 donors) exposed to mouse-derived myelin in combination with and without medium exposed to KNS42 (NM-HGG) tumor cells were profiled by bulk RNA-sequencing. Heatmaps showing Z-scored expression of DESEQ2 normalized counts of genes of interest. **E,** Representative Visium spatial plots color-coded for gene expression of SPP1, GPNMB, CD68, CD8A, PLIN2, and a MES-like signature score. Visium data acquired from Ren et al^65^. **F,** Heatmaps visualizing the average log fold change of cytokines and chemokines upregulated and downregulated in SPP1-high spots as compared to other spots. Visualized genes were commonly upregulated or downregulated amongst all five DMG Visium sections.

To validate the relevance of these genes, we used published spatial transcriptomic data of five diagnostic DMG sections generated using 10X Visium^65^ and scored each spot for the expression of *SPP1*, *GPNMB*, *CD68*, *CD8A* or MES-like signature genes^24^. We observed the co-occurrence of *GPNMB* and *SPP1* with spots that scored high for the MES-like signature as visualized in Figure 4E and Supplementary Figure 5A, in line with our CosMx observations. Interestingly, the spots with the highest MES-like signature score showed lower gene expression of *MBP* (myelin basic protein) and *PLP1* (proteolipid protein 1), which encode the predominant myelin proteins (Supplementary Fig. 5B), underscoring the role of myelin within these regions.

Next, we looked at the cytokines and chemokines upregulated within the *SPP1*-high spots. We identified 20 cytokines and chemokines that were commonly upregulated in *SPP1*- high spots in all five DMG samples analyzed (Fig. 4F). We identified immunosuppressive genes that we previously found to be upregulated in the myelin- and TCM-exposed macrophages *in vitro* (*CXCL16*, *CXCL8*, *TNFSF14, EBI3, GRN,* among others). Other upregulated genes include *IL1RN (*a decoy receptor that can inhibit pro-inflammatory signaling by binding IL1β)^66^ and *IL18* (a driver of CD8^+^ T cell exhaustion)^67^. Moreover, the chemokines and cytokines *CCL20*, *CCL28*, *VEGF*, and *EGF* are known promoters of angiogenesis and EMT^68,69,70^. Downregulated genes in *SPP1*-high spots include the pro-inflammatory cytokine *IL17D*^71^, and genes from the Semaphorin family (*SEMA6C, SEMA6D, SEMA4D),* known for their role in axonal guidance and cytokine release^72^.

Collectively, our data present SPP1^+^/GPNMB^+^ MES-like myeloid cells as a new and prominent component of the pHGG’s TIME. We demonstrated that the combination of myelin uptake with TCM recapitulates the transcriptomic profile observed in regions with elevated *SPP1*/*GPNMB* expression and shows that an interplay between the tumor cells and myeloid cells contributes to the establishment of immunosuppressive environments.

### SPP1^+^ myeloid cells are associated with reduced CD8^+^ T cell proximity

Our data suggests that the chemokines and cytokines secreted by SPP1^+^/GPNMB^+^ myeloid cells contribute to the formation of an immunosuppressive tumor microenvironment (Fig. 4). Moreover, given the role of SPP1 in immune evasive mechanisms in other cancers^28^, we hypothesized that MES-like myeloid cells would affect the influx of T cells in pHGG. cIF revealed clear areas with increased numbers of CD8^+^ T cells in both DMG and NM-HGG (Fig. 5A). Notably, these areas also contained high numbers of CD163^+^ myeloid cells, but were devoid of SPP1^+^ myeloid cells, as visualized in density plots (Fig. 5B). We confirmed our findings in a larger patient cohort by performing cIF on a sequential section of the TMA (Fig. 1). Similarly, this analysis revealed CD8^+^ T cells situated at a greater distance from SPP1^+^/GPNMB^+^ myeloid cells (Fig. 5C, Supplementary fig. 6A). We quantified cell-cell interactions by neighborhood permutation testing and observed that CD8^+^ T cells are more likely to interact with other myeloid cells rather than with SPP1^+^/GPNMB^+^ myeloid cells (Fig. 5D). Correspondingly, in four out of five DMG sections in the Visium dataset we observed the distance of *CD8A*-high spots to the nearest *SPP1-*high spot to be significantly higher than their distance to the nearest *CD68*-high spot (Supplementary Fig. 6B), reconfirming our protein level observations.

**Figure 5.**
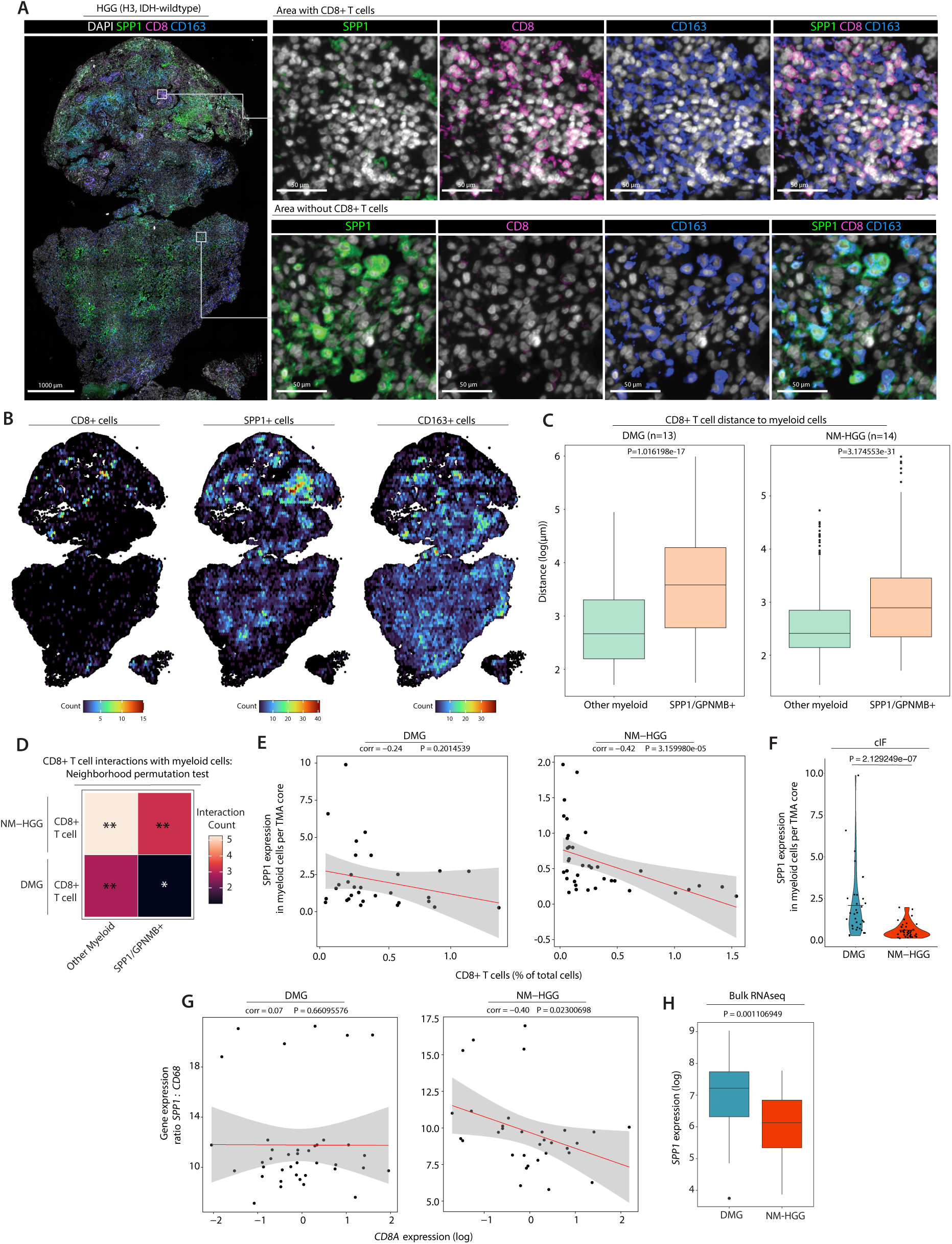
SPP1^+^ myeloid cells are associated with reduced CD8^+^ T cell proximity. A, Representative immunofluorescence image of a NM-HGG sample showing protein expression of SPP1, CD8, CD163. Magnifications of a region with CD8^+^ T cells, and without CD8^+^ T cells. **B,** Density plot showing the count of CD8^+^ cells, SPP1^+^ cells, and CD163^+^ cells detected within the tissue of (**A**). Cells were binned in a 10 μm grid to enhance visibility. **C,** Box plots comparing the distance of CD8^+^ T cells to the nearest other myeloid cell as determined by cIF on the TMA consisting of samples of 13 DMG patients and 14 NM-HGG patients. Dots represent outliers. P-values were determined by the Mann-Whitney U test. **D,** Heatmap visualizing the spatial cellular co- localization and interaction as determined by neighborhood permutation testing within the imcRtools package. A significance threshold of P=0.05 was used. * = 0.004995, ** = 0.0009990. **E,** Scatter plots showing the correlation between the expression of SPP1 in myeloid cells per TMA core and the CD8^+^ T cell count as a percentage of total cells detected. Each dot represents a TMA core (DMG n=34, NM-HGG n=37). P-values and correlation determined by Spearman correlation testing. **F,** Violin plot comparing the SPP1 expression in myeloid cells in each TMA core between DMG patients (n=34) and NM-HGG patients (n=37). Each dot represents a TMA core. P-values were determined by Mann-Whitney U test. **G,** Scatter plots showing the correlation between the gene expression ratio SPP1: CD68, reflecting the SPP1 expression in myeloid cells, and CD8A expression in bulk RNA-seq data of DMG patients (n=39) and NM-HGG patients (n=31). Each dot represents a patient. P-values and correlation determined by Spearman correlation testing.

Next, we hypothesized that tumors and/or tumor regions with higher expression of SPP1 in myeloid cells would contain fewer CD8^+^ T cells. Indeed, we observed a statistically significant inverse correlation between SPP1 expression in myeloid cells and the percentage of CD8^+^ T cells in NM-HGGs (Fig. 5E). Interestingly, DMG myeloid cells expressed more SPP1 compared to NM-HGGs (Fig. 5F), providing a potential explanation for the overall lower percentage of CD8^+^ T cells in DMG compared to NM-HGGs (Fig. 1F). To further support our findings at the protein level, we asked whether similar correlations could be observed in our bulk RNA-seq data of pHGG patients. In this analysis, we identified a significant inverse correlation between the *SPP1:CD68* ratio, reflecting *SPP1* expression in myeloid cells, and the *CD8A* expression in NM-HGGs (Fig. 5G). Again, we see significantly higher expression of *SPP1* in DMGs compared to NM-HGGs (Fig. 5H), highlighting a possible midline-specific response of myeloid cells under pathological conditions, as has been shown previously for midbrain-specific microglia^73^. We next acquired bulk RNA-seq data from the OpenPBTA^74^, which contains annotated CNS tumor locations for each sample. This allowed us to compare *SPP1* expression between midline and hemispheric tumors. Similar to our cIF and bulk RNA- seq data, we observe significantly higher expression of *SPP1* in tumors arising in the midline (Supplementary Fig. 6C). Furthermore, we observed a significant increase in *GPNMB* and MES-like signature expression at recurrence or autopsy for the NM-HGGs but not for DMGs (Fig. 6D), perhaps reflecting different progression-free survival times and/or treatment regimens for these two different pHGG entities.

Thus, our spatial analyses show a negative spatial correlation at the cellular level between SPP1^+^ (MES-like) myeloid cells and CD8^+^ T cells, as well as at a regional (single TMA core) level and in bulk RNA-seq data of pHGG. These findings are supported by studies indicating the immunosuppressive roles of SPP1 in other cancers^28,29^ and its role as a potent T cell suppressor.

## Discussion

The spatial architecture of tumors has recently emerged as an additional dimension in better understanding the TIME, the communication between individual cells, and predicting response to immunotherapies^17,15^. With the ultimate aim of improving future (immuno)therapies for pHGG, we here provide a comprehensive overview of the TIME within a broad set of pHGG tumor samples collected at diagnosis. Despite some differences in the composition of the TIME between different clinical entities, which may be at least partly, explained by tumor location ^5,75^, the overall number of T cells in all pHGGs was observed to be very low. Since the efficacy of most immunotherapies depends on CD8^+^ T cell influx, we aimed to get a better understanding of the low number of T cells in pHGGs. Therefore, we decided to focus on myeloid cells, the largest compartment of the pHGG’s TIME, with known effect on T cell recruitment, and studied the possible interplay between pHGG myeloid cells and T cells.

Using single-cell spatial transcriptomics, we identified a large population of SPP1^+^/GPNMB^+^ myeloid cells in both DMG and NM-HGG, indicating that this may be a conserved and dominant cell type in pediatric brain tumors. Indeed, Liu *et al.* and DeSisto *et al.* recently also reported the expression of MES-like genes in DMG-associated myeloid cells, although predominantly in adult patients^24,76^. *GPNMB* and *SPP1* expression has been associated with lipid-related metabolic^77^ and demyelination disorders^27^. Indeed, we found increased expression of lipid metabolism-related genes in SPP1^+^/GPNMB^+^ myeloid cells compared to other myeloid cells. The acquisition of this so-called lipid-laden state has been shown to be driven by lipid uptake from the microenvironment under pathological conditions^36^, and may originate from the conserved developmental mechanism of synaptic engulfment by MG, for which SPP1 is required^78,27^. In line with this, we show that the uptake of lipids in the form of myelin, a lipid-containing structure abundant in the CNS, contributes to the lipid-laden phenotype of myeloid cells. How SPP1^+^/GPNMB^+^ myeloid cells acquire their lipid-laden phenotype remains to be explored. Hypoxia likely plays a major role as it has been shown to induce both MES-like tumor cells^79^ and lipid accumulation in macrophages^80^. Furthermore, hypoxia-induced factors have been associated with macrophage migration, which may explain why macrophages were predominantly detected in MES-like tumor cell-enriched regions within the DMG spatial transcriptomic data.

Spatially, SPP1^+^/GPNMB^+^ myeloid cells were negatively correlated with CD8^+^ T cells, which may be due to the lack of T cell-recruiting chemokines. Indeed, SPP1^+^/GPNMB^+^ regions were characterized by the upregulated expression of immunosuppressive cytokines and chemokines. Interestingly, tumor-associated CD163^+^ myeloid cells, classically defined as an immunosuppressive population, were found close to CD8^+^ T cells. This suggests the existence of different myeloid cell populations with different immunosuppressive properties. Where SPP1^+^/GPNMB^+^ myeloid cells may exclude CD8^+^ T cells from tumors, CD163^+^ myeloid cells may exert their immunosuppressive action differently, for example by expressing immune checkpoints that were found to positively correlate with CD163 expression (such as PD-L1, PD-1, TIM-3, LAG-3)^81,82^. Thus, our data suggest that multiple immunosuppressive myeloid populations must be targeted for optimal therapeutic efficacy.

In summary, our study provides novel insights into the interaction between myeloid cells, T cells and tumor cells within the TIME of pHGG. We identify SPP1 and GPNMB as highly upregulated in these tumors, potentially contributing to the limited efficacy of chimeric antigen receptor T cell (CAR-T) therapies observed in clinical settings^2,4^. Our findings suggest that cellular immunotherapies may be more effective when combined with strategies targeting these immunosuppressive myeloid cells. Future studies should even more so focus on characterizing the myeloid compartment and its spatial organization in treated pHGGs. As early phase clinical trials predominantly involve treated and recurrent patients, understanding the TIME after initial treatment is crucial. We hypothesize that SPP1^+^/GPNMB^+^ myeloid cells become even more dominant post-treatment, particularly following radiotherapy, which preferentially eliminates well-oxygenated cells, while sparing those in hypoxic regions. If these myeloid cells represent a major component of irradiated pHGGs, co-targeting these populations could enhance treatment efficacy.

## Methods

### TMA generation

We collected archived formalin-fixed paraffin-embedded (FFPE) tumor material from the tissue biobank at the Princess Máxima Center in Utrecht, The Netherlands. Samples were obtained as patients underwent surgical resection. Regions of interest were identified by an experienced neuropathologist based on diagnostic hematoxylin and eosin sections. Three regions were selected per tissue of which cylindrical punches of 1-mm diameter were taken. These punches were combined into a tissue micro-array (TMA) of which 4-μm thick sections were generated for cyclical immunofluorescence imaging and spatial molecular imaging.

### Antigen retrieval

FFPE tissue sections were deparaffinized by washing in xylene (3 x 3 min) followed by rehydration in a series of graded alcohol for 1 min each (2 x 100%, 2 x 95%, 1x 70%). After washing (2 x 1 min) in deionized water, the sections were put in Target Retrieval Solution, pH 9 (Agilent Dako). Antigens were retrieved for 40 min at 95°C. After 1 hour at room temperature, the sections were washed for 5 min in deionized water.

### Cyclical immunofluorescence imaging

cIF was applied as previously described^83^. After antigen retrieval, a barrier was drawn surrounding the tissue using a hydrophobic pen. Tissue was exposed to a blocking solution consisting of 100 mM NH4Cl (ThermoFisher), 150 mM maleimide (Merck), and 10% donkey serum (Merck) in PBS for 1 hour in a humidified chamber at RT. Afterwards, the blocking solution was replaced with a primary antibody staining solution containing 100 mM NH4Cl and 5% donkey serum. The primary antibodies used were CD31 (R&D Systems, AF3628), CD45 (LSBio, LS-B14248-300), TMEM119 (Cell Signaling, 41134S), IBA1 (Fujifilm Wako, 019- 19741), CD68 (Abcam, ab955), CD163 (ThermoFisher, MA5-11458), CD206 (R&D Systems, MAB25341), FOXP3 (Thermofisher, MA5-16365), CD8 (Bio-Rad, MCA1817T), CD4 (abcam, ab133616), PLIN2 (Biotechne, NB110-40877SS), GPNMB (R&D Systems, AF2550-SP), and SPP1 (R&D Systems, AF1433-SP). Sections were incubated for 2.5 h in a humidified chamber on an orbital shaker followed by washing in PBS (3 x 5 min). Sections were exposed to a secondary antibody staining solution (containing 100 mM NH4Cl, 5% donkey serum, and DAPI (Biolegend)) for 1 h at RT in the dark. Secondary antibodies used were Anti-Rabbit Cy5 (Jackson ImmunoResearch, 711-175-152), Anti-Mouse AF555 (ThermoFisher, A-31570), Anti-Goat AF488 (ThermoFisher, A-11055), all raised in donkey. After washing (3 x 5 min) in PBS, SlowFade Gold antifade mounting medium (Invitrogen) was applied on the sections and a coverslip was mounted. Imaging of the slides was performed on a Leica DMi8 Thunder imaging system using a HC PL APO 20x/0.80 objective. After imaging, the coverslips were removed in PBS followed by washing in PBS (3 x 5 min). Antibody removal was performed by applying elution buffer (Lunaphore) or a chemical antibody removal buffer (0.5 M glycine, 3 M guanidium chloride, 2 M urea, 40 mM tris(2-carboxyethyl)phosphine, ultrapure water) to the sections for 3 min. Sections were washed in PBS (3 x 5 min) which allowed for the cIF cycle to start again by applying the blocking buffer.

### Image processing and spatial analysis

Images were merged using Leica LASX Software. Image alignment for each cycle of cIF was based on the DAPI signal for which we used HiFiAlignmentTool (https://github.com/jhausserlab/HiFiAlignmentTool). The script was adapted to facilitate LIF files. Sometimes images from different cycles require alignment beyond linear adjustments (Supplementary Fig. 1A), therefore we developed a more advanced tool available at https://github.com/Dream3DLab/CycFluoCoreg. Initially, the tool applies an affine (linear) transformation to each channel to align the images to a DAPI reference, ensuring overall centering. This is followed by a B-spline (non-linear) transformation, which refines the alignment by correcting local deformations, resulting in better overlap. In addition, CycFluoCoreg optimizes the use of computational resources to balance processing speed and accuracy. The final output is an aligned composite image that is ready for further analysis.

Composite images were imported into QuPath (v0.4.4)^84^ for analysis. Each section was annotated followed by nuclear detection and segmentation using a cell expansion of 2.5 µm. For each marker, an object classifier using RandomTrees was trained. These separate object classifiers for each marker were then combined to create a composite classifier which was applied for cell identification. Spatial coordinates, distances, and marker intensities were exported for downstream analysis in R using the imcRtools package^23^. We applied the same image analysis to cIF images of full tissue sections. To generate spatial density maps of detected cell types, we binned cells within 10 µm to generate a grid. The density map with cell count was plotted on the tissue outline.

### CosMx tissue processing, image acquisition, cell segmentation

TMA sections were dispatched to NanoString Technologies for processing. CosMx SMI was performed as previously described^85^. Briefly, overnight drying of the slides at 37°C was followed by deparaffinization, antigen retrieval and proteinase-mediated permeabilization to allow for probe penetration. Antigen retrieval was carried out at 100°C for 15 min in Leica ER1 buffer and proteinase permeabilization was performed using 3.0 μg/mL proteinase K in 1x PBS for 30 min at 40°C. RNA in situ hybridization (ISH) was conducted using 1 nM RNA-ISH probes, which were applied to the sections and incubated overnight at 60°C. Washing steps were performed to remove non-specifically bound probes and a flow cell was assembled on top of the slide to enable cyclic RNA readout on the CosMx using a 16-digit encoding strategy. Following completion of all cycles, a cocktail of fluorescently-labeled protein markers and a nuclear marker were applied to the samples to collect morphology images that allowed for cell segmentation. The morphology marker cocktail consisted of GFAP, CD298+B2M+18s rRNA, and Histone H3. DAPI was used as the nuclear marker.

For imaging, twenty-five 0.5 mm x 0.5 mm fields of view (FOVs) were placed at the center of each TMA core that fitted within the CosMx scanning area. The CosMx optical system, equipped with a custom water immersion objective (x22.77 NA 1.1) and widefield illumination from a mix of lasers and LEDs (385 nm, 488 nm, 530 nm, 590 nm, and 656 nm), was utilized for image acquisition. Images were captured using a ATX204S-MC camera, which incorporates an IMX531 Sony industrial CMOS sensor. Image processing involved registration, feature extraction, localization, and decoding of individual RNA transcripts, followed by cell segmentation using a machine learning algorithm based on Cellpose, as described previously^85^. Based on transcript location, the segmentation allowed for mapping of the transcripts in the registered images to the corresponding cell.

The applied probe set consisted of the 1000-plex RNA panel for Universal Cell Typing and a custom designed 30-plex probe set of glioblastoma-based targets that can be used for Cell-Cell Interaction studies.

### CosMx cell type prediction

We separated the DMG samples from the NM-HGGs for further analysis. Scanpy^33^ and Squidpy^86^ were used to analyze all CosMx data. Cells with less than 80 genes detected were excluded from further analysis as well as FOVs with poor quality. Data was normalized and log transformed followed by principal component analysis using 20 components combined with default parameters. To remove batch effects, we applied the Harmony algorithm^87^. A neighborhood graph was computed using scanpy.pp.neighbors before clusters were determined by Leiden clustering. We used the Tangram package as described previously^88^ to predict cell types from a published scRNA-sequencing dataset to the DMG sample subset^24^. Liu *et al.* scRNA-sequencing data was downloaded from GEO (GSE184357). For the NM- HGGs we went through several iterations of manual annotation based on marker gene expression within the computed Leiden clusters.

### CosMx spatial analysis

As each field of view (FOV) within the CosMx data captured a different TMA core, we calculated neighborhood enrichment z-scores for each FOV separately using squidpy.gr.nhood_enrichment. Results for each FOV were aggregated and aligned to generate neighborhood enrichment heatmaps. Spatial distances between cell types were determined for each FOV separately using squidpy.tl.var_by_distance. Following this, the distances were combined for all FOVs to generate a complete dataset which could be imported into R for visualization.

### Ligand-receptor analysis

FOVs were labeled as MES-like or AC-like/OPC-like based on the fraction of MES-like tumor cells. Macrophages and microglia, as well as MES-like tumor cells were separated from the dataset to perform ligand-receptor (LR) analysis for these regions specifically. The Liana method was used to predict LR interactions^89^. We used liana.mt.rank_aggregate to aggregate and integrate predictions from separate methods to analyze LR interactions. Consensus interactions were visualized.

### Flow cytometry and imaging

Peripheral blood was obtained from healthy donors with approval from the Ethical Committee of the University Medical Center Utrecht (UMCU), The Netherlands. Monocytes were isolated from peripheral blood using a Pan Monocyte Isolation kit (Miltenyi Biotec) following manufacturer’s instructions. Monocytes were cultured for 6 days in RPMI1640 (Gibco) supplemented with 10% fetal bovine serum (FBS) and 50 ng/ml M-CSF (Biolegend) to induce differentiation to naïve macrophages. KNS-42 cells (H3G34-mutant) were obtained from the Japanese Cancer Research Resources Bank (JCRB) and cultured in RPMI1640 supplemented with 10% FBS. All cells were cultured at 37 °C and 5% CO2 and a humidity of 95%. Tumor-conditioned medium (TCM) was collected after being exposed to KNS-42 cells for 72 hours. Naïve macrophages were re-seeded in RPMI1640, or TCM. Medium was supplemented with 1 µg/ml M-CSF. After 24h, the cells were exposed to medium containing 1 mg/ml of myelin for 3 h. Part of the cells were harvested and shipped to Novogene for RNA-sequencing or fixed in the culture plate for 15 min with 4% paraformaldehyde for confocal imaging. The cells were stained for GPNMB (R&D Systems, AF2550-SP) and DAPI using the same reagents and procedure as described in section *Cyclical immunofluorescence imaging.* Confocal imaging of the cells was performed on a Zeiss LSM880 using a x20 objective. After myelin exposure, the other part of the cells were resuspended in Human TruStain FcX (Biolegend) in staining buffer (PBS with 2% FBS and 2 mM EDTA) for 10 minutes to initiate flow cytometric analysis. After washing, cells were resuspended in LIVE/DEAD Near IR (876) dye (Invitrogen) in staining buffer. After 30 min at RT, cells were stained with CD14-BV785 (Biolegend), CD163-BV605 (Biolegend), GPNMB-PE (Invitrogen), BODIPY 493/503 (Thermofisher). Flow cytometry was performed on a CytoFLEX LX (Beckman Coulter). FlowJo 10.10.0 was used for analysis.

### Bulk RNA-sequencing of patient samples

Total RNA was isolated from fresh frozen tumor material using the AllPrep DNA/RNA/Protein Mini Kit (QIAGEN). RNA-seq libraries were generated with 300ng of RNA using the KAPA RNA HyperPrep Kit with RiboErase (Roche) and sequenced on a NovaSeq 6000 system (2 × 150 bp) (Illumina). RNA sequencing data were processed according to the GATK 4.0 best practices workflow for variant calling using a wdl- and cromwell-based workflow^90^. Raw sequencing reads were aligned using Star (version 2.7.0f) ^91^ to GRCh38 and Gencode version 31 ^92^. featureCounts from Rsubread (v1.32.4) ^93^ was used to extract gene expression counts.

Raw counts were normalized to counts per million (CPM). Normalized counts were log-transformed to calculate Z-scores. To determine cell type signature scores based on^16,24^, the average Z-scores of the genes were calculated per sample.

### Bulk RNA-sequencing of cultured cells

HSJD-DIPG-007 (H3K27-altered), HSJD-DIPG-012 (H3K27-altered), HSJD-GBM-001 (H3 wildtype), HSJD-GBM-002 (H3G34-mutant) cells were obtained from Sant Joan de Déu Barcelona Hospital. M354AAB-TO cells, derived from a 7-year old patient with a H3K27- altered pediatric type diffuse midline glioma, were obtained from the Princess Máxima Center for Pediatric Oncology. HSJD-DIPG-007, HSJD-DIPG-012, and M354AAB-TO cells were grown in tumor stem medium (TSM) consisting of Neurobasal-A and DMEM/F-12 (1:1) medium supplemented with 10 mM HEPES buffer, 1 x MEM non-essential amino acids, 1% GlutaMAX, 1 mM Sodium pyruvate,1 x B-27 minus vitamin-A (all Thermofisher), 10 ng/ml PDGF-AA, 10 ng/ml PDGF-BB, 20 ng/ml bFGF, 20 ng/ml EGF (all Peprotech), 2 μg/ml Heparin (StemCell Technologies), and for M354AAB-TO also supplemented with 0.2% Cultrex Reduced Growth Factor Basement Membrane Extract (Biotechne) and 1 x N-2 supplement (Thermofisher). All cells were cultured at 37 °C and 5% CO2 and a humidity of 95%. DMG (HSJD-DIPG-007, HSJD-DIPG-012, M354AAB-TO) and NM-HGG (HSJD-GBM-001, HSJD-GBM-002, KNS-42) cells were exposed to 5 μg/ml recombinant human SPP1 protein (abcam) for 24 hours. Cells were snap frozen and shipped to Novogene for RNA sequencing. RNA integrity was assessed using the Bioanalyzer 2100 system (Agilent Technologies). Non strand-specific library preparation was performed by Novogene followed by sequencing on the Illumina NovaSeq X Plus platform. Reads were aligned to the reference genome using HISAT2 (v2.0.5) ^94^. featureCounts was used to extract gene expression counts. DESeq2 ^95^ was used to normalize counts and compare expression between treatment groups.

### Publicly available Visium data

We acquired raw Visium spatial transcriptomic data of five DMG sections published by Ren *et al.,* (2023) from GEO (GSE194329)^65^. All Visium analyses were performed using Scanpy. We normalized and log transformed the data before determining the highly variable genes. MES-like signature was based on Liu *et al*. Spots enriched in *SPP1*, *CD68*, or *CD8* were identified for each section by computing the 90^th^ percentile using NumPy. Coordinates of these spots were extracted and cdist from scipy.spatial was used to compute the spatial distances between spots. Mann-Whitney U test was used for comparison of spatial distances to *CD8* between two groups. Differential gene expression analysis focused on cytokines and chemokines was performed between *SPP1*-high spots and other spots. Cytokine and chemokine lists were acquired via ImmPort^96^.

### Publicly available bulk RNA-sequencing data

Processed bulk RNA-sequencing data from the Open Pediatric Brain Tumor Atlas (OpenPBTA, v23) was downloaded from PedcBioPortal^74^. Derived cell lines were excluded. Samples were excluded based on “Cancer Type” and “Detailed Cancer Type”.

### Statistics and reproducibility

Sample size was limited by availability of diagnostic material within the tissue biobank and was therefore not determined by statistical test. Python (3.9.6), R version 4.3.2 (2023-10-31), and GraphPad Prism Version 9.5.1 were used for all statistical analyses.

## Data and Code availability

The spatial transcriptomic data (NanoString CosMx) used in this project is available at https://zenodo.org/records/14225941. The bulk RNA-sequencing data of patients is access-restricted and access can be requested at the European Genome-phenome Archive under the accession number EGAS00001008002. Published DMG single cell RNA-sequencing data were downloaded under accession code GSE184357. Published DMG Visium data were downloaded under accession code GSE194329. Published bulk RNA-sequencing data was downloaded from https://pedcbioportal.org/.

The image alignment tool is available at https://github.com/Dream3DLab/CycFluoCoReg. No other custom code was created or implemented for the analyses in this study. Standard workflows, along with open-source R and Python packages were utilized as presented in Methods and Supplementary Methods.

## Author contributions

T.J.M.B., R.H., M.E.G.K., J.L., D.P.S., H.G.S., A.C.R., D.G.V., and A.Z. conceived and designed the experiments and analyses. M.E.G.K., R.H., E.W.H., F.C.A.R., M.W. performed tissue acquisition and processing. T.J.M.B., J.A.S.L., A.L.K., Y.S., and A.Z. performed experiments. T.J.M.B., R.H., J.A.S.L. analyzed the data with contributions from V.O.T., D.P.S., and C.R.M., L.A. provided technical advice and reagents. R.L.I. and M.M. generated the image alignment tool. H.G.S., A.C.R., D.G.V. and A.Z. supervised the project. T.J.M.B., D.G.V., and A.Z. wrote the paper. R.H., J.A.S.L., M.G.R., J.I.B., D.P.S., and A.C.R. edited the paper. All authors read and approved the paper.

## Competing interests

The authors declare no competing interests. A.Z. currently works at Genmab, The Netherlands.

### Acknowledgements

We thank the Imaging Center of the Princess Máxima Center for their excellent support. We thank the Flow Cytometry core facility of the Princess Máxima Center. We thank NanoString Technologies Inc. for performing the CosMx experiment. We thank D. Castigliego of the University Medical Center Utrecht for generating the tissue micro-array. Patient samples were acquired from the Princess Máxima Center Biobank. Healthy donor blood was acquired from the Mini Donor Service of the University Medical Center Utrecht.

## Supplementary Figures

**Supplementary figure 1.**
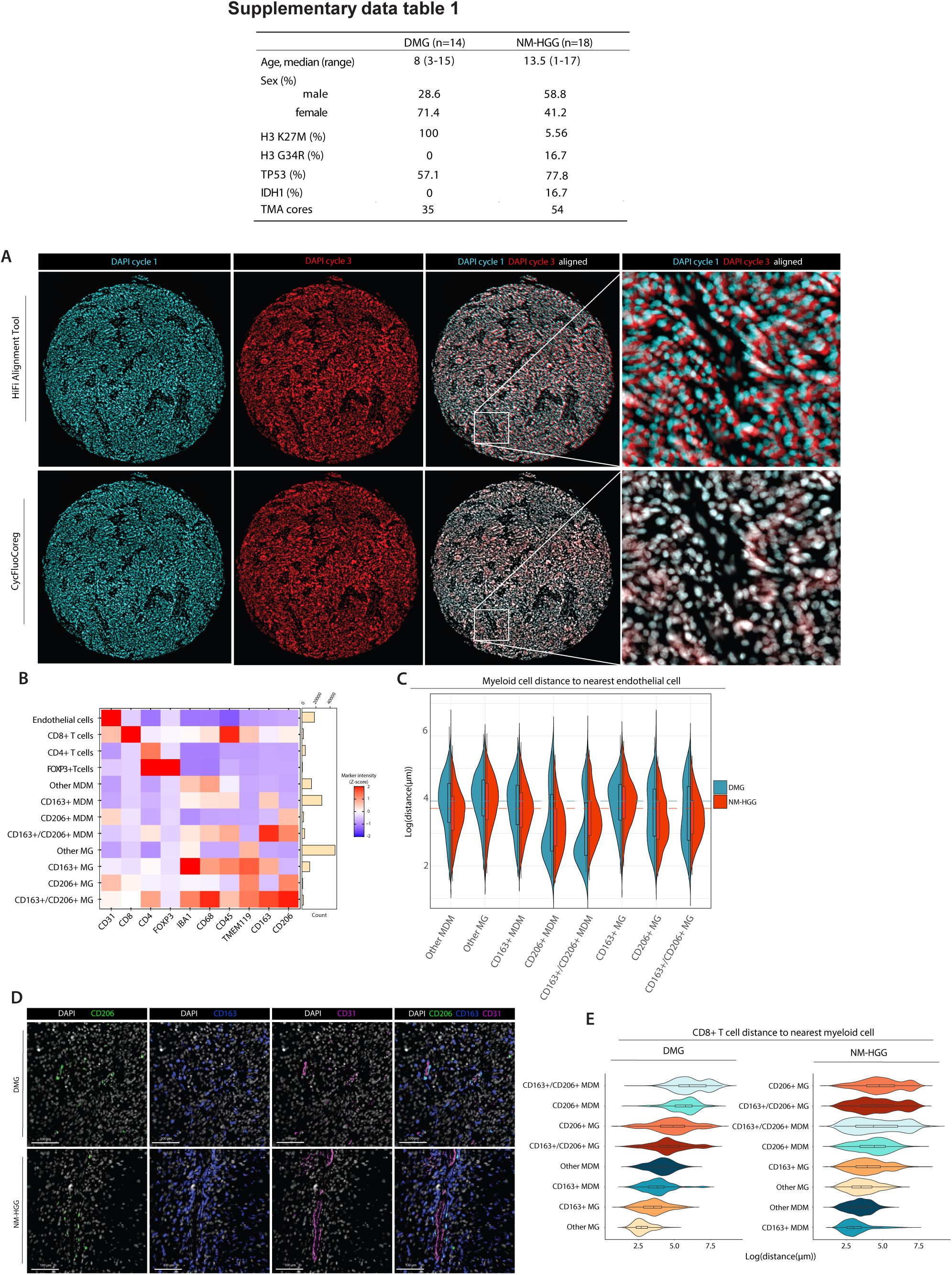
Alignment, marker expression and single-cell distance analysis of cIF data. A, Representative immunofluorescence images generated using cIF of a TMA core. The same images of three cIF cycles were aligned using the HiFi Alignment Tool^83^ and CycFluoCoReg. DAPI signal imaged during the first and the third cycles of cIF visualized. **B,** Heatmap visualization of the Z-scored mean fluorescence intensity of each marker detected per identified cell type in the total cIF dataset (DMG and NM-HGG combined). Horizontal bar plot shows the count of each identified cell type. **B,** Violin plots visualizing the distance of myeloid cell types to the nearest endothelial cell. Dashed lines indicate the mean distance for DMG and for NM-HGG in blue and red, respectively. **C,** Representative immunofluorescence images of CD206, CD163, and CD31. **D,** Violin plots visualizing the distance of CD8+ T cells to the nearest myeloid cell for DMG and NM-HGG.

**Supplementary figure 2.**
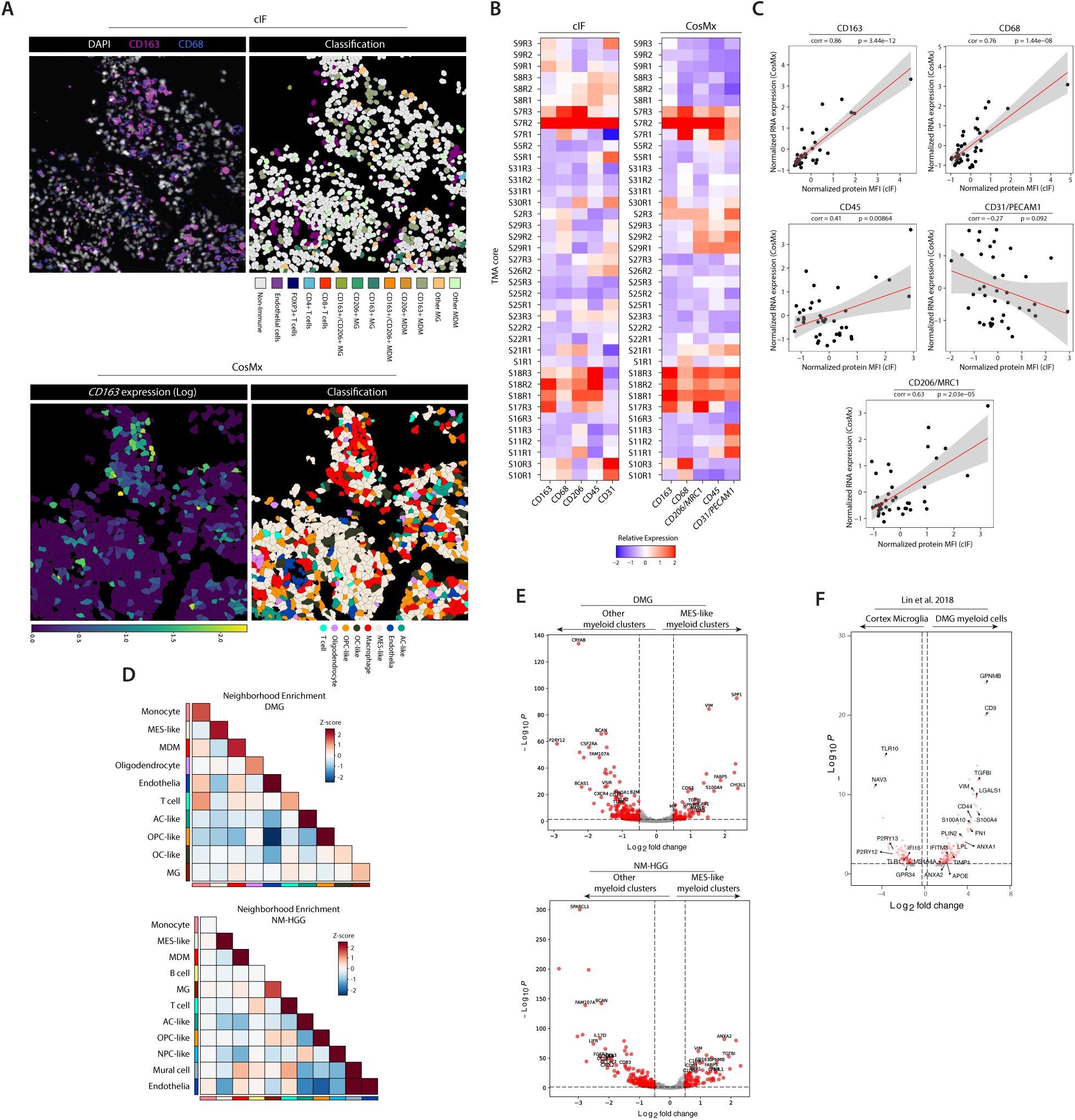
Cellular classification and analysis of CosMx data. A, Representative immunofluorescence image and its subsequent classification of cell types, generated by cIF of one TMA core. RNA expression of CD163 as captured by CosMx of the same TMA core, and classification of cell types detected. **B,** Heatmap visualizing the relative expression of proteins detected using cIF and the corresponding RNA expression detected by NanoString CosMx in the same samples of an adjacent slice of the TMA. **C,** Scatter plots showing the correlation of RNA expression and protein expression detected using CosMx and cIF, respectively. P-values and correlation determined by Spearman correlation testing. **D,** Heatmap visualization of neighborhood analysis performed per FOV captured by CosMx. **E,** Differentially expressed genes between MES-like myeloid clusters and other myeloid clusters. **D,** Differentially expressed genes between DMG myeloid cells and healthy cortex microglia. Data from Lin et al. (2018) ^97^.

**Supplementary figure 3.**
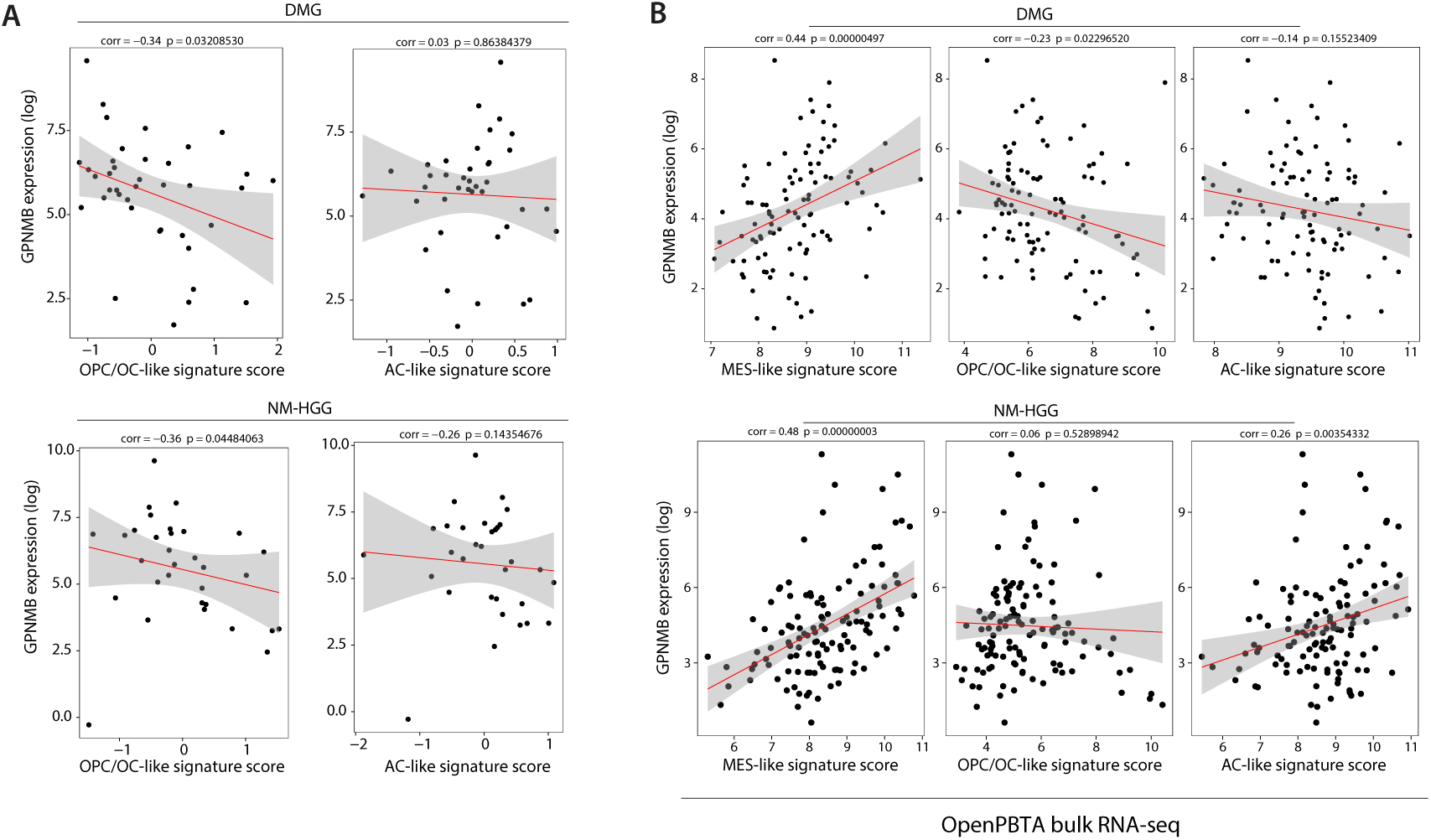
Correlation between GPNMB expression and tumor state signature scores in bulk RNA-seq data. A, Scatter plots showing the correlation between GPNMB expression and tumor state signature scores in an in-house cohort of 39 DMG patients and 31 NM-HGG patients and in (**B**) OpenPBTA bulk RNA-seq data ^74^. Each dot represents one patient. P-values and correlation determined by Spearman correlation testing.

**Supplementary figure 4.**
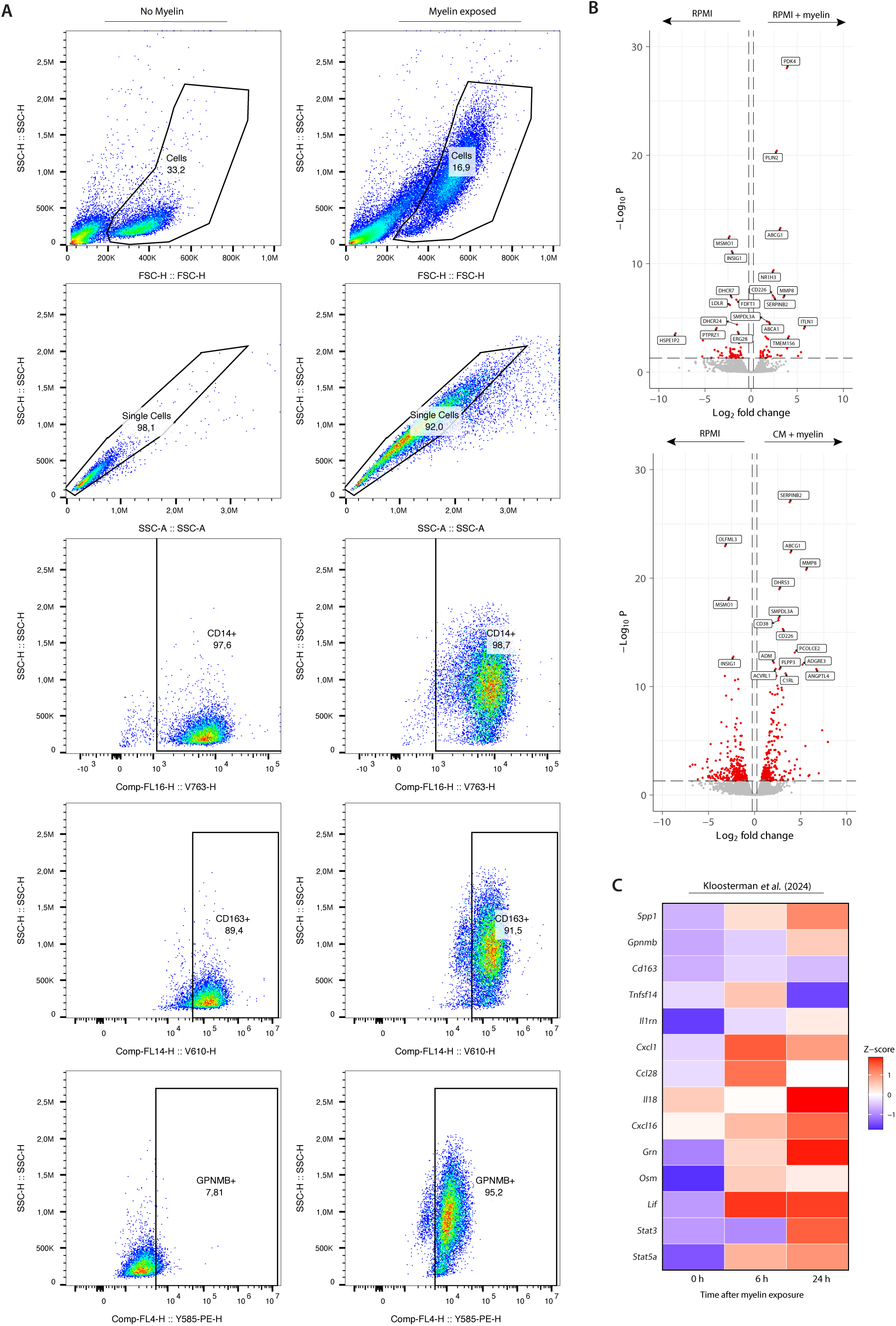
Flow cytometry gating strategy and bulk RNA sequencing data of in vitro generated lipid-laden macrophages. **A**, Cells were gated on single cells followed by CD14^+^ cells. The left panel shows representative plots for a non-myelin exposed sample while the right panel shows representative plots for a myelin-exposed sample. **B**, Volcano plot showing the differentially expressed genes between in vitro generated naïve macrophages and myelin-exposed macrophages, or naïve macrophages and macrophages exposed to myelin and tumor-conditioned medium. **C,** Heatmap visualizing the (Z-scored) expression of genes of interest determined by bulk RNA sequencing of mouse-derived macrophages before (0 h) and 6 h or 24 h after exposure to myelin debris. n=3 per timepoint. Data acquired from Kloosterman et al.^32^

**Supplementary figure 5.**
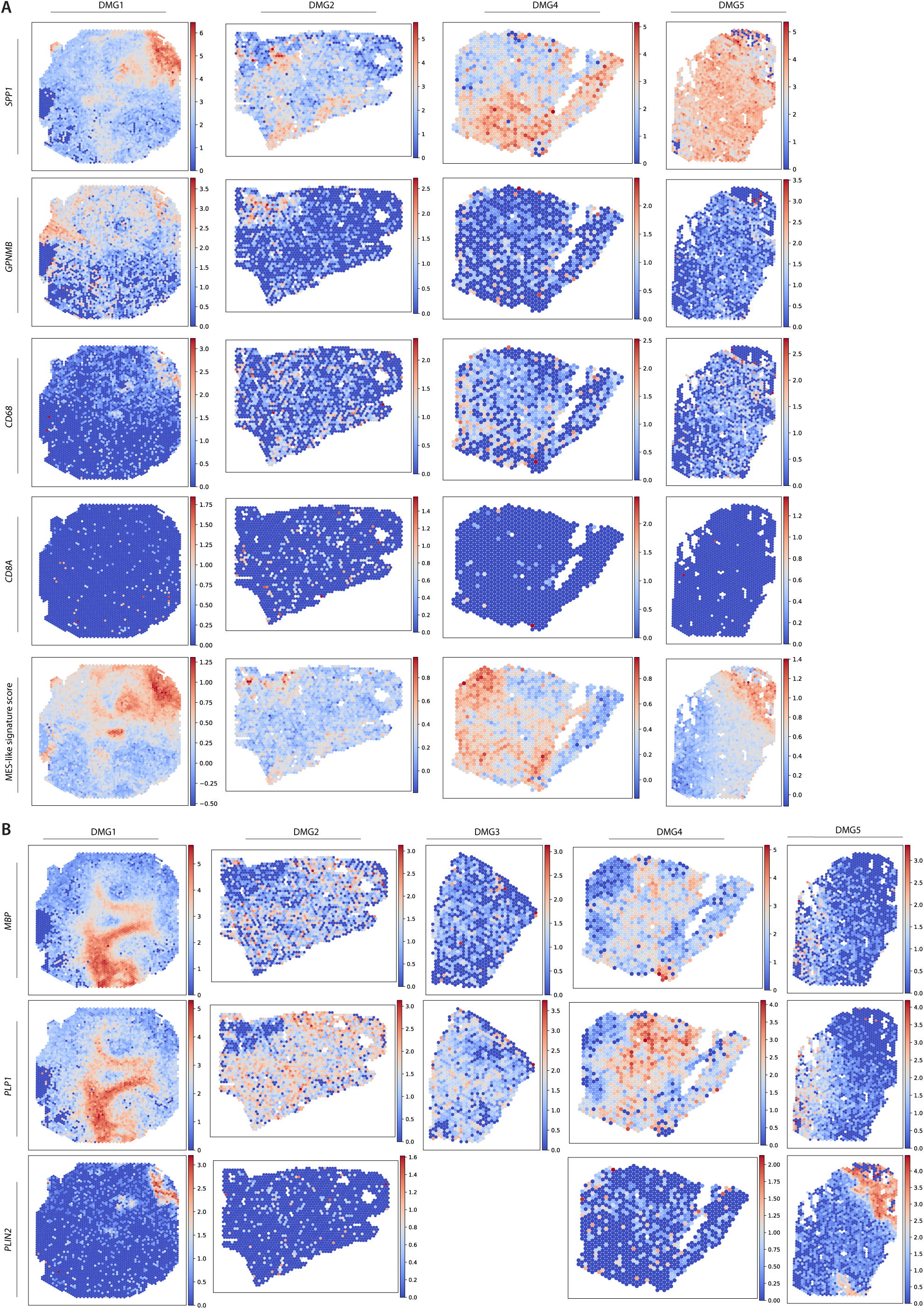
Visium spatial plots. A, Visium spatial plots showing spatial expression of genes SPP1, GPNMB, CD68, CD8A, and a MES-like signature score. Data generated by Ren et al. (2023)^65^. **B**, Visium spatial plots of genes MBP, PLP1, PLIN2.

**Supplementary figure 6.**
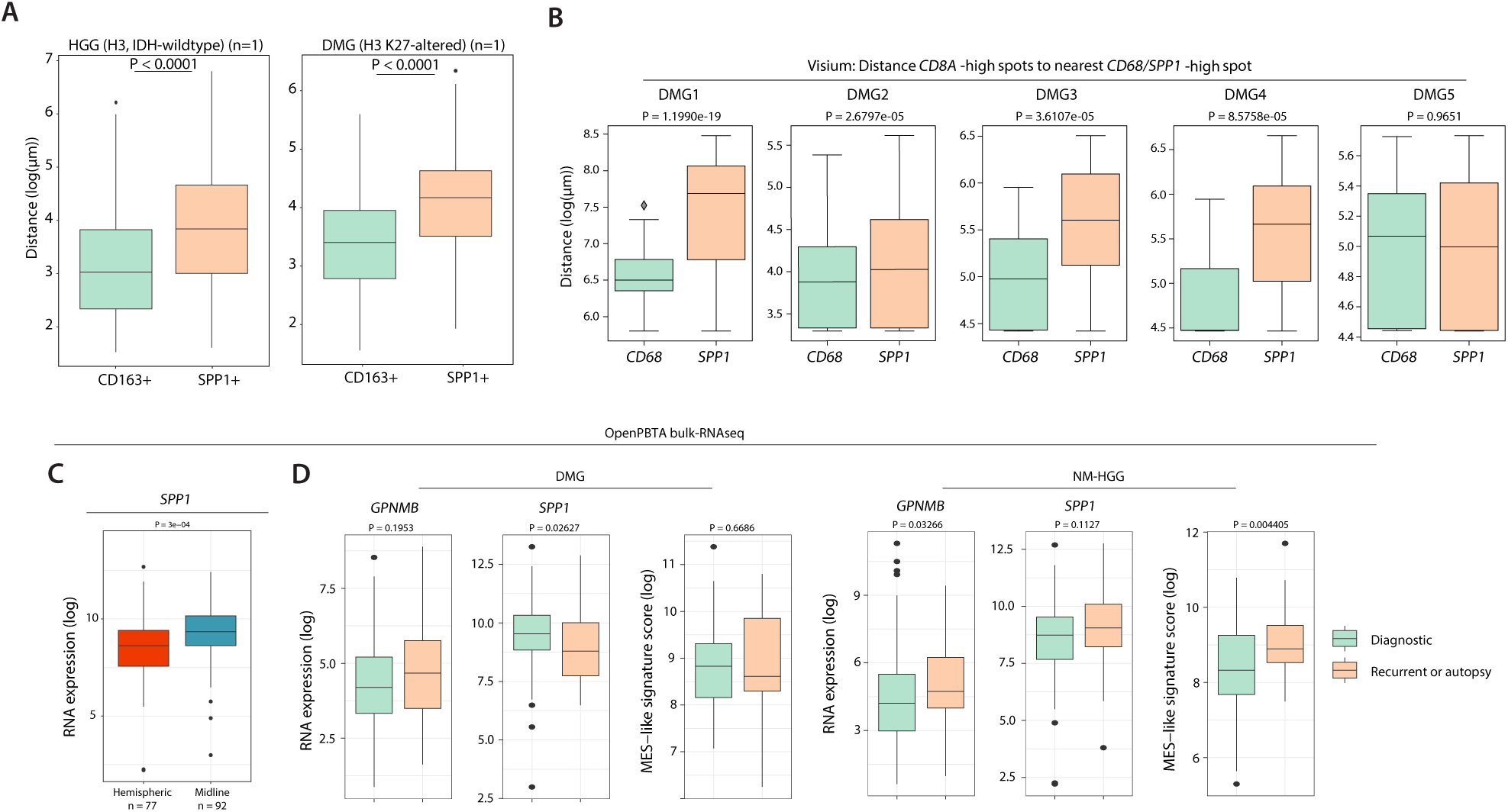
Distance analysis of SPP1+/GPNMB+ myeloid cells and OpenPBTA analysis. A, Box plots comparing the distance of CD8+ T cells to the nearest myeloid cell expressing either CD163 or SPP1 in a full NM-HGG section and a full DMG section. Dots represent outliers. P-values were determined by the Mann-Whitney U test. **B,** Box plots comparing the distance of spots with high CD8 expression to the nearest spots with either high CD68 expression or SPP1 expression as detected in five DMG Visium sections. P-values were determined by the Mann-Whitney U test. Diamonds represent outliers. **C,** Box plot comparing the expression of SPP1 between tumors diagnosed in hemispheric structures and midline structures. P-value determined by Mann-Withney U test. **D,** Box plots comparing the expression of GPNMB, SPP1, and a MES-like signature score between samples obtained at diagnosis and samples obtained at recurrence or during autopsy. Dots represent outliers. P-values determined by Mann-Withney U test. Data obtained from the OpenPBTA^74^.

**Supplementary figure 7.**
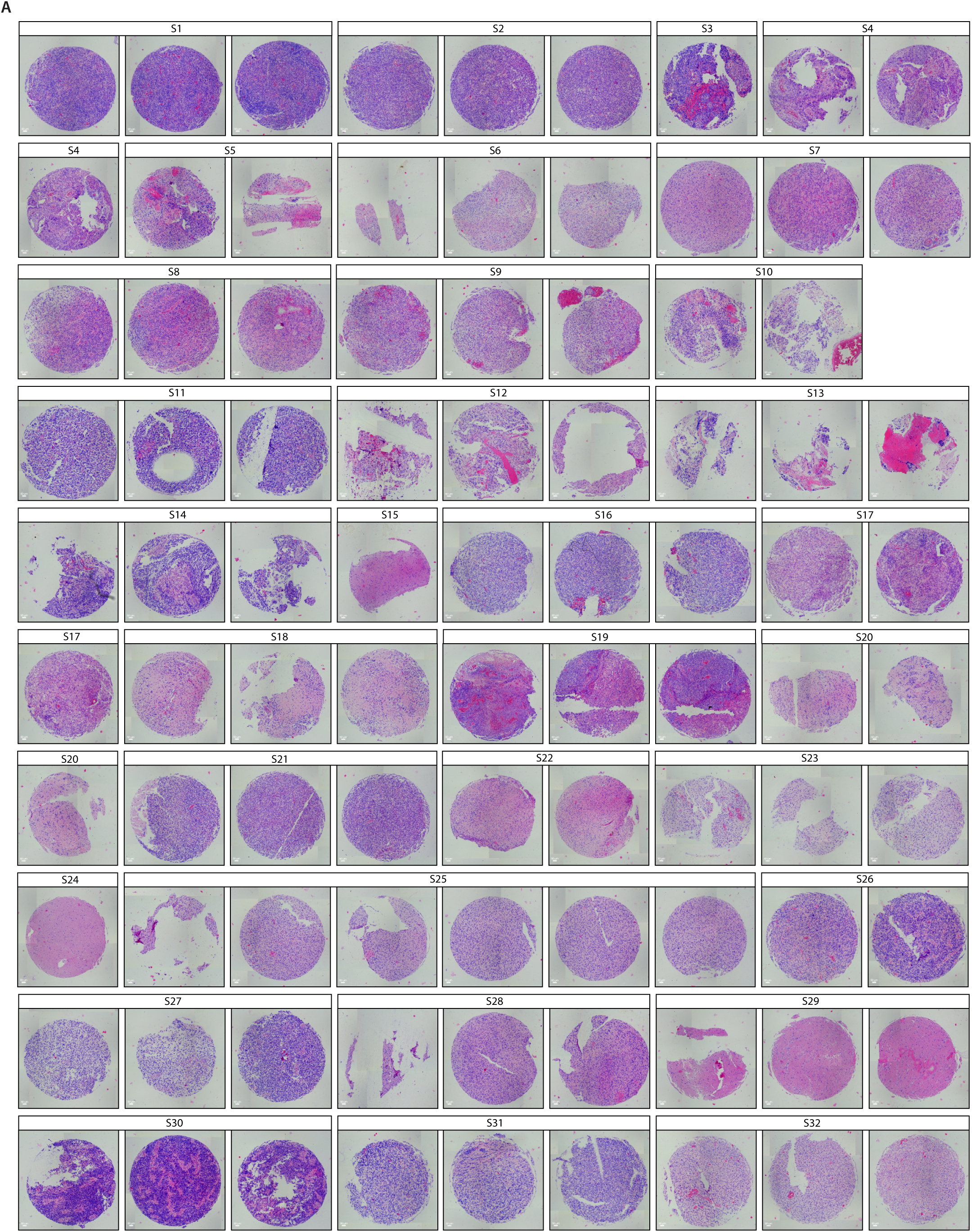
**A**, Hematoxylin and eosin staining of TMA cores after full cIF run.

## References

1. Mackay A, Burford A, Carvalho D, et al. Integrated Molecular Meta-Analysis of 1,000 Pediatric High-Grade and Diffuse Intrinsic Pontine Glioma. Cancer Cell. 2017;32:520–537. doi:10.1016/j.ccell.2017.08.017

2. Majzner RG, Ramakrishna S, Yeom KW, et al. GD2-CAR T cell therapy for H3K27M-mutated diffuse midline gliomas. Nature. 2022;603. doi:10.1038/s41586-022-04489-4

3. Gállego Pérez-Larraya J, Garcia-Moure M, Labiano S, et al. Oncolytic DNX-2401 Virus for Pediatric Diffuse Intrinsic Pontine Glioma. New England Journal of Medicine. 2022;386(26):2471-2481. doi:10.1056/NEJMoa2202028

4. Vitanza NA, Wilson AL, Huang W, et al. Intraventricular B7-H3 CAR T Cells for Diffuse Intrinsic Pontine Glioma: Preliminary First-in-Human Bioactivity and Safety. Cancer Discov. 2023;13(1):114–131. doi:10.1158/2159-8290.CD-22-0750

5. Grabovska Y, Mackay A, O’Hare P, et al. Pediatric pan-central nervous system tumor analysis of immune-cell infiltration identifies correlates of antitumor immunity. Nature Communications 2020 11:*1*. 2020;11(1):1-15. doi:10.1038/s41467-020-18070-y

6. Mackay A, Burford A, Molinari V, et al. Molecular, Pathological, Radiological, and Immune Profiling of Non-brainstem Pediatric High-Grade Glioma from the HERBY Phase II Randomized Trial. Cancer Cell. 2018;33(5):829–842.e5. doi:10.1016/j.ccell.2018.04.004

7. Lieberman NAP, Degolier K, Kovar HM, et al. Characterization of the immune microenvironment of diffuse intrinsic pontine glioma: implications for development of immunotherapy. Neuro Oncol. 2019;21(1):83–94. doi:10.1093/NEUONC/NOY145

8. Petralia F, Tignor N, Reva B, et al. Integrated Proteogenomic Characterization across Major Histological Types of Pediatric Brain Cancer. Cell. 2020;183(7):1962–1985.e31. doi:10.1016/j.cell.2020.10.044

9. Ausejo-Mauleon I, Labiano S, de la Nava D, et al. TIM-3 blockade in diffuse intrinsic pontine glioma models promotes tumor regression and antitumor immune memory. Cancer Cell. 2023;41(11):1911–1926.e8. doi:10.1016/j.ccell.2023.09.001

10. Mass E, Nimmerjahn F, Kierdorf K, Schlitzer A. Tissue-specific macrophages: how they develop and choreograph tissue biology. Nat Rev Immunol. 2023;23(9):1. doi:10.1038/S41577-023-00848-Y

11. Mosser DM, Edwards JP. Exploring the full spectrum of macrophage activation. Nat Rev Immunol. 2008;8(12):958. doi:10.1038/NRI2448

12. Lazarov T, Juarez-Carreño S, Cox N, Geissmann F. Physiology and diseases of tissue-resident macrophages. Nature 2023 618:7966. 2023;618(7966):698-707. doi:10.1038/S41586-023-06002-X

13. DeNardo DG, Brennan DJ, Rexhepaj E, et al. Leukocyte complexity predicts breast cancer survival and functionally regulates response to chemotherapy. Cancer Discov. 2011;1(1):54–67. doi:10.1158/2159-8274.CD-10-0028/42904/

14. Seo YD, Jiang X, Sullivan KM, et al. Mobilization of CD8+ T cells via CXCR4 blockade facilitates PD-1 checkpoint therapy in human pancreatic cancer. Clinical Cancer Research. 2019;25(13):3934–3945. doi:10.1158/1078-0432.CCR-19-0081/74932/AM/MOBILIZATION-OF-CD8-T-CELLS-VIA-CXCR4-BLOCKADE

15. Piyadasa H, Angelo M, Bendall SC. Spatial proteomics of tumor microenvironments reveal why location matters. Nature Immunology 2023 24:*4*. 2023;24(4):565-566. doi:10.1038/s41590-023-01471-8

16. Zheng Y, Carrillo-Perez F, Pizurica M, Heiland DH, Gevaert O. Spatial cellular architecture predicts prognosis in glioblastoma. Nature Communications 2023 14:*1*. 2023;14(1):1-16. doi:10.1038/s41467-023-39933-0

17. Wang XQ, Danenberg E, Huang CS, et al. Spatial predictors of immunotherapy response in triple-negative breast cancer. Nature 2023 621:7980. 2023;621(7980):868-876. doi:10.1038/s41586-023-06498-3

18. Gil-Jimenez A, van Dijk N, Vos JL, et al. Spatial relationships in the urothelial and head and neck tumor microenvironment predict response to combination immune checkpoint inhibitors. Nature Communications 2024 15:*1*. 2024;15(1):1-15. doi:10.1038/s41467-024-46450-1

19. Mantovani A, Marchesi F, Malesci A, Laghi L, Allavena P. Tumour-associated macrophages as treatment targets in oncology. Nature Reviews Clinical Oncology 2017 14:*7*. 2017;14(7):399- 416. doi:10.1038/nrclinonc.2016.217

20. Masuda T, Amann L, Monaco G, et al. Specification of CNS macrophage subsets occurs postnatally in defined niches. Nature 2022 604:7907. 2022;604(7907):740-748. doi:10.1038/s41586-022-04596-2

21. Sankowski R, Süß P, Benkendorff A, et al. Multiomic spatial landscape of innate immune cells at human central nervous system borders. Nature Medicine 2023 30:*1*. 2023;30(1):186-198. doi:10.1038/s41591-023-02673-1

22. Louis DN, Perry A, Wesseling P, et al. The 2021 WHO Classification of Tumors of the Central Nervous System: a summary. Neuro Oncol. 2021;23(8):1231-1251. doi:10.1093/NEUONC/NOAB106

23. Windhager J, Zanotelli VRT, Schulz D, et al. An end-to-end workflow for multiplexed image processing and analysis. Nature Protocols 2023 18:*11*. 2023;18(11):3565-3613. doi:10.1038/s41596-023-00881-0

24. Liu I, Jiang L, Samuelsson ER, et al. The landscape of tumor cell states and spatial organization in H3-K27M mutant diffuse midline glioma across age and location. Nature Genetics 2022 54:*12*. 2022;54(12):1881-1894. doi:10.1038/s41588-022-01236-3

25. Jessa S, Mohammadnia A, Harutyunyan AS, et al. K27M in canonical and noncanonical H3 variants occurs in distinct oligodendroglial cell lineages in brain midline gliomas. Nat Genet. 2022;54(12):1865–1880. doi:10.1038/S41588-022-01205-W

26. Neftel C, Laffy J, Filbin MG, et al. An Integrative Model of Cellular States, Plasticity, and Genetics for Glioblastoma. Cell. 2019;178(4):835–849.e21. doi:10.1016/J.CELL.2019.06.024

27. Lan Y, Zhang X, Liu S, et al. Fate mapping of Spp1 expression reveals age-dependent plasticity of disease-associated microglia-like cells after brain injury. Immunity. 2024;57(2):349–363.e9. doi:10.1016/j.immuni.2024.01.008

28. Bill R, Wirapati P, Messemaker M, et al. *CXCL9:SPP1* macrophage polarity identifies a network of cellular programs that control human cancers. Science (1979). 2023;381(6657):515–524. doi:10.1126/science.ade2292

29. Qi J, Sun H, Zhang Y, et al. Single-cell and spatial analysis reveal interaction of FAP+ fibroblasts and SPP1+ macrophages in colorectal cancer. Nat Commun. 2022;13(1). doi:10.1038/S41467-022-29366-6

30. Xiong A, Zhang J, Chen Y, Zhang Y, Yang F. Integrated single-cell transcriptomic analyses reveal that GPNMB-high macrophages promote PN-MES transition and impede T cell activation in GBM. EBioMedicine. 2022;83:104239. doi:10.1016/j.ebiom.2022.104239

31. Yalcin F, Haneke H, Efe IE, et al. Tumor associated microglia/macrophages utilize GPNMB to promote tumor growth and alter immune cell infiltration in glioma. Acta Neuropathol Commun. 2024;12(1):1–21. doi:10.1186/S40478-024-01754-7/FIGURES/8

32. Kloosterman DJ, Erbani J, Boon M, et al. Macrophage-mediated myelin recycling fuels brain cancer malignancy. Cell. 2024;187(19):5336–5356.e30. doi:10.1016/J.CELL.2024.07.030

33. Wolf FA, Angerer P, Theis FJ. SCANPY: Large-scale single-cell gene expression data analysis.Genome Biol. 2018;19(1):1–5. doi:10.1186/S13059-017-1382-0/FIGURES/1

34. Chen AX, Gartrell RD, Zhao J, et al. Single-cell characterization of macrophages in glioblastoma reveals MARCO as a mesenchymal pro-tumor marker. Genome Med. 2021;13(1):1–13. doi:10.1186/S13073-021-00906-X/FIGURES/5

35. Abdelfattah N, Kumar P, Wang C, et al. Single-cell analysis of human glioma and immune cells identifies S100A4 as an immunotherapy target. Nature Communications 2022 13:*1*. 2022;13(1):1-18. doi:10.1038/s41467-022-28372-y

36. Masetti M, Carriero R, Portale F, et al. Lipid-loaded tumor-associated macrophages sustain tumor growth and invasiveness in prostate cancer. J Exp Med. 2022;219(2). doi:10.1084/JEM.20210564

37. Yang M, Zhang G, Wang Y, et al. Tumour-associated neutrophils orchestrate intratumoural IL- 8-driven immune evasion through Jagged2 activation in ovarian cancer. Br J Cancer. 2020;123(9):1404. doi:10.1038/S41416-020-1026-0

38. Pascual-García M, Bonfill-Teixidor E, Planas-Rigol E, et al. LIF regulates CXCL9 in tumor-associated macrophages and prevents CD8+ T cell tumor-infiltration impairing anti-PD1 therapy. Nat Commun. 2019;10(1). doi:10.1038/S41467-019-10369-9

39. Justilien V, Regala RP, Tseng IC, et al. Matrix Metalloproteinase-10 Is Required for Lung Cancer Stem Cell Maintenance, Tumor Initiation and Metastatic Potential. PLoS One. 2012;7(4):e35040. doi:10.1371/JOURNAL.PONE.0035040

40. Sarkar S, Nuttall RK, Liu S, Edwards DR, Yong VW. Tenascin-C stimulates glioma cell invasion through matrix metalloproteinase-12. Cancer Res. 2006;66(24):11771–11780. doi:10.1158/0008-5472.CAN-05-0470

41. Heid HW, Moll R, Schwetlick I, Rackwitz HR, Keenan TW. Adipophilin is a specific marker of lipid accumulation in diverse cell types and diseases. Cell Tissue Res. 1998;294(2):309–321. doi:10.1007/S004410051181/METRICS

42. Haney MS, Pálovics R, Munson CN, et al. APOE4/4 is linked to damaging lipid droplets in Alzheimer’s disease microglia. Nature 2024 628:8006. 2024;628(8006):154-161. doi:10.1038/s41586-024-07185-7

43. Grajchen E, Wouters E, Van De Haterd B, et al. CD36-mediated uptake of myelin debris by macrophages and microglia reduces neuroinflammation. J Neuroinflammation. 2020;17(1):1–14. doi:10.1186/S12974-020-01899-X/FIGURES/6

44. Loix M, Wouters E, Vanherle S, et al. Perilipin-2 limits remyelination by preventing lipid droplet degradation. Cellular and Molecular Life Sciences. 2022;79(10):1–16. doi:10.1007/S00018-022-04547-0/METRICS

45. Luo M, Qiu Z, Tang X, et al. Inhibiting Cyclin B1-treated Pontine Infarction by Suppressing Proliferation of SPP1+ Microglia. Mol Neurobiol. 2023;60(4):1782–1796. doi:10.1007/S12035-022-03183-W/FIGURES/6

46. Wolf KJ, Shukla P, Springer K, et al. A mode of cell adhesion and migration facilitated by CD44- dependent microtentacles. Proc Natl Acad Sci U S A. 2020;117(21):11432–11443. doi:10.1073/PNAS.1914294117/SUPPL_FILE/PNAS.1914294117.SM03.MOV

47. Hara T, Chanoch-Myers R, Mathewson ND, et al. Interactions between cancer cells and immune cells drive transitions to mesenchymal-like states in glioblastoma. Cancer Cell. 2021;39(6):779–792.e11. doi:10.1016/j.ccell.2021.05.002

48. Desgrosellier JS, Cheresh DA. Integrins in cancer: biological implications and therapeutic opportunities. Nature Reviews Cancer 2010 10:*1*. 2010;10(1):9-22. doi:10.1038/NRC2748

49. D’Angelo RC, Liu XW, Najy AJ, et al. TIMP-1 via TWIST1 induces EMT phenotypes in human breast epithelial cells. Molecular Cancer Research. 2014;12(9):1324–1333. doi:10.1158/1541-7786.MCR-14-0105/80724/AM/TIMP-1-VIA-TWIST1-INDUCES-EMT-PHENOTYPES-IN-HUMAN

50. Chen L, Cai S, Wang J mei, Huai Y ying, Lu PH, Chu Q. BRDT promotes ovarian cancer cell growth. Cell Death & Disease 2020 11:*11*. 2020;11(11):1-14. doi:10.1038/s41419-020-03225-y

51. Karimbayli J, Pellarin I, Belletti B, Baldassarre G. Insights into the structural and functional activities of forgotten Kinases: PCTAIREs CDKs. Molecular Cancer 2024 23:*1*. 2024;23(1):1-20. doi:10.1186/S12943-024-02043-6

52. Cain RJ, Ridley AJ. Phosphoinositide 3-kinases in cell migration. Biol Cell. 2009;101(1):13–29. doi:10.1042/BC20080079

53. Yi Y, Tsai SH, Cheng JC, et al. APELA promotes tumour growth and cell migration in ovarian cancer in a p53-dependent manner. Gynecol Oncol. 2017;147(3):663–671. doi:10.1016/J.YGYNO.2017.10.016

54. Aguadé-Gorgorió J, Jami-Alahmadi Y, Calvanese V, et al. MYCT1 controls environmental sensing in human haematopoietic stem cells. Nature 2024 630:8016. 2024;630(8016):412-420. doi:10.1038/s41586-024-07478-x

55. Shih CH, Chang YJ, Huang WC, et al. EZH2-mediated upregulation of ROS1 oncogene promotes oral cancer metastasis. Oncogene 2017 36:*47*. 2017;36(47):6542-6554. doi:10.1038/onc.2017.262

56. Tan XY, Li YT, Li HH, et al. WNT2–SOX4 positive feedback loop promotes chemoresistance and tumorigenesis by inducing stem-cell like properties in gastric cancer. Oncogene 2023 42:*41*. 2023;42(41):3062-3074. doi:10.1038/s41388-023-02816-1

57. Nian Z, Dou Y, Shen Y, et al. Interleukin-34-orchestrated tumor-associated macrophage reprogramming is required for tumor immune escape driven by p53 inactivation. Immunity. 2024;57(10):2344–2361.e7. doi:10.1016/j.immuni.2024.08.015

58. Kloosterman DJ, Farber M, Boon M, Erbani J, Akkari L. Protocol for studying macrophage lipid crosstalk with murine tumor cells. STAR Protoc. 2024;5(4). doi:10.1016/J.XPRO.2024.103421

59. Kim MJ, Sun HJ, Song YS, et al. CXCL16 positively correlated with M2-macrophage infiltration, enhanced angiogenesis, and poor prognosis in thyroid cancer. Scientific Reports 2019 9:*1*. 2019;9(1):1-10. doi:10.1038/s41598-019-49613-z

60. Wu Y, Zhan S, Chen L, et al. TNFSF14/LIGHT promotes cardiac fibrosis and atrial fibrillation vulnerability via PI3Kγ/SGK1 pathway-dependent M2 macrophage polarisation. J Transl Med. 2023;21(1):1–18. doi:10.1186/S12967-023-04381-3/FIGURES/9

61. Hildenbrand K, Bohnacker S, Menon PR, et al. Human interleukin-12α and EBI3 are cytokines with anti-inflammatory functions. Sci Adv. 2023;9(43). doi:10.1126/SCIADV.ADG6874

62. Cheung PF, Yang JJ, Fang R, et al. Progranulin mediates immune evasion of pancreatic ductal adenocarcinoma through regulation of MHCI expression. Nature Communications 2022 13:*1*. 2022;13(1):1-18. doi:10.1038/s41467-021-27088-9

63. Agresta L, Lehn M, Lampe K, et al. CD244 represents a new therapeutic target in head and neck squamous cell carcinoma. J Immunother Cancer. 2020;8(1). doi:10.1136/JITC-2019-000245

64. Shouval DS, Biswas A, Goettel JA, et al. Interleukin-10 receptor signaling in innate immune cells regulates mucosal immune tolerance and anti-inflammatory macrophage function. Immunity. 2014;40(5):706–719. doi:10.1016/J.IMMUNI.2014.03.011

65. Ren Y, Huang Z, Zhou L, et al. Spatial transcriptomics reveals niche-specific enrichment and vulnerabilities of radial glial stem-like cells in malignant gliomas. Nature Communications 2023 14:*1*. 2023;14(1):1-19. doi:10.1038/s41467-023-36707-6

66. Chen L, Huang H, Zheng X, et al. IL1R2 increases regulatory T cell population in the tumor microenvironment by enhancing MHC-II expression on cancer-associated fibroblasts. J Immunother Cancer. 2022;10(9):e004585. doi:10.1136/JITC-2022-004585

67. Lutz V, Hellmund VM, Picard FSR, et al. IL18 Receptor Signaling Regulates Tumor-Reactive CD8+ T-cell Exhaustion via Activation of the IL2/STAT5/mTOR Pathway in a Pancreatic Cancer Model. Cancer Immunol Res. 2023;11(4):421–434. doi:10.1158/2326-6066.CIR-22-0398/716498/AM/IL-18-RECEPTOR-SIGNALING-REGULATES-TUMOR-REACTIVE

68. Cheng XS, Li YF, Tan J, et al. CCL20 and CXCL8 synergize to promote progression and poor survival outcome in patients with colorectal cancer by collaborative induction of the epithelial–mesenchymal transition. Cancer Lett. 2014;348(1-2):77–87. doi:10.1016/J.CANLET.2014.03.008

69. Huang G, Tao L, Shen S, Chen L. Hypoxia induced CCL28 promotes angiogenesis in lung adenocarcinoma by targeting CCR3 on endothelial cells. Sci Rep. 2016;6. doi:10.1038/SREP27152

70. De Palma M, Biziato D, Petrova T V. Microenvironmental regulation of tumour angiogenesis. Nature Reviews Cancer 2017 17:*8*. 2017;17(8):457-474. doi:10.1038/NRC.2017.51

71. McGeachy MJ, Cua DJ, Gaffen SL. The IL-17 Family of Cytokines in Health and Disease. Immunity. 2019;50(4):892–906. doi:10.1016/J.IMMUNI.2019.03.021

72. Zhou Y, Gunput RAF, Pasterkamp RJ. Semaphorin signaling: progress made and promises ahead. Trends Biochem Sci. 2008;33(4):161–170. doi:10.1016/J.TIBS.2008.01.006

73. Abellanas MA, Zamarbide M, Basurco L, et al. Midbrain microglia mediate a specific immunosuppressive response under inflammatory conditions. J Neuroinflammation. 2019;16(1):1–15. doi:10.1186/S12974-019-1628-8/FIGURES/7

74. Shapiro JA, Gaonkar KS, Spielman SJ, et al. OpenPBTA: The Open Pediatric Brain Tumor Atlas. Cell Genomics. 2023;3(7). doi:10.1016/J.XGEN.2023.100340

75. Bailey CP, Wang R, Figueroa M, Zhang S, Wang L, Chandra J. Computational immune infiltration analysis of pediatric high-grade gliomas (pHGGs) reveals differences in immunosuppression and prognosis by tumor location. Comput Syst Oncol. 2021;1(3). doi:10.1002/CSO2.1016

76. DeSisto J, Donson AM, Griesinger AM, et al. Tumor and immune cell types interact to produce heterogeneous phenotypes of pediatric high-grade glioma. Neuro Oncol. 2024;26(3):538–552. doi:10.1093/NEUONC/NOAD207

77. Prabata A, Ikeda K, Rahardini EP, Hirata KI, Emoto N. GPNMB plays a protective role against obesity-related metabolic disorders by reducing macrophage inflammatory capacity. Journal of Biological Chemistry. 2021;297(5). doi:10.1016/j.jbc.2021.101232

78. De Schepper S, Ge JZ, Crowley G, et al. Perivascular cells induce microglial phagocytic states and synaptic engulfment via SPP1 in mouse models of Alzheimer’s disease. Nature Neuroscience 2023 26:*3*. 2023;26(3):406-415. doi:10.1038/s41593-023-01257-z

79. Ravi VM, Will P, Kueckelhaus J, et al. Spatially resolved multi-omics deciphers bidirectional tumor-host interdependence in glioblastoma. Cancer Cell. 2022;40(6):639–655.e13. doi:10.1016/J.CCELL.2022.05.009/ATTACHMENT/DB6FC941-11CD-492C-97BE-4AB2A596AE8F/MMC3.XLSX

80. Bensaad K, Favaro E, Lewis CA, et al. Fatty acid uptake and lipid storage induced by HIF-1α contribute to cell growth and survival after hypoxia-reoxygenation. Cell Rep. 2014;9(1):349–365. doi:10.1016/j.celrep.2014.08.056

81. Liu S, Zhang C, Maimela NR, et al. Molecular and clinical characterization of CD163 expression via large-scale analysis in glioma. Oncoimmunology. 2019;8(7). doi:10.1080/2162402X.2019.1601478

82. Zhang H, Zhang N, Wu W, et al. Pericyte mediates the infiltration, migration, and polarization of macrophages by CD163/MCAM axis in glioblastoma. iScience. 2022;25(9). doi:10.1016/J.ISCI.2022.104918

83. Watson SS, Duc B, Kang Z, et al. Microenvironmental reorganization in brain tumors following radiotherapy and recurrence revealed by hyperplexed immunofluorescence imaging. Nature Communications 2024 15:*1*. 2024;15(1):1-16. doi:10.1038/s41467-024-47185-9

84. Bankhead P, Loughrey MB, Fernández JA, et al. QuPath: Open source software for digital pathology image analysis. Scientific Reports 2017 7:*1*. 2017;7(1):1-7. doi:10.1038/s41598-017-17204-5

85. He S, Bhatt R, Brown C, et al. High-plex imaging of RNA and proteins at subcellular resolution in fixed tissue by spatial molecular imaging. Nature Biotechnology 2022 40:*12*. 2022;40(12):1794-1806. doi:10.1038/s41587-022-01483-z

86. Palla G, Spitzer H, Klein M, et al. Squidpy: a scalable framework for spatial omics analysis.Nature Methods 2022 19:*2*. 2022;19(2):171-178. doi:10.1038/s41592-021-01358-2

87. Korsunsky I, Millard N, Fan J, et al. Fast, sensitive and accurate integration of single-cell data with Harmony. Nature Methods 2019 16:*12*. 2019;16(12):1289-1296. doi:10.1038/s41592-019-0619-0

88. Biancalani T, Scalia G, Buffoni L, et al. Deep learning and alignment of spatially resolved single-cell transcriptomes with Tangram. Nature Methods 2021 18:*11*. 2021;18(11):1352- 1362. doi:10.1038/s41592-021-01264-7

89. Dimitrov D, Türei D, Garrido-Rodriguez M, et al. Comparison of methods and resources for cell-cell communication inference from single-cell RNA-Seq data. Nat Commun. Published online 2022. doi:10.1038/s41467-022-30755-0

90. McKenna A, Hanna M, Banks E, et al. The Genome Analysis Toolkit: A MapReduce framework for analyzing next-generation DNA sequencing data. Genome Res. 2010;20(9):1297–1303. doi:10.1101/GR.107524.110

91. Dobin A, Davis CA, Schlesinger F, et al. STAR: ultrafast universal RNA-seq aligner. Bioinformatics. 2013;29(1):15–21. doi:10.1093/BIOINFORMATICS/BTS635

92. Frankish A, Diekhans M, Jungreis I, et al. GENCODE 2021. Nucleic Acids Res. 2021;49(D1):D916-D923. doi:10.1093/NAR/GKAA1087

93. Liao Y, Smyth GK, Shi W. The R package Rsubread is easier, faster, cheaper and better for alignment and quantification of RNA sequencing reads. Nucleic Acids Res. 2019;47(8):e47–e47. doi:10.1093/NAR/GKZ114

94. Kim D, Paggi JM, Park C, Bennett C, Salzberg SL. Graph-based genome alignment and genotyping with HISAT2 and HISAT-genotype. Nature Biotechnology 2019 37:*8*. 2019;37(8):907-915. doi:10.1038/s41587-019-0201-4

95. Love MI, Huber W, Anders S. Moderated estimation of fold change and dispersion for RNA-seq data with DESeq2. Genome Biol. 2014;15(12):1–21. doi:10.1186/S13059-014-0550-8/FIGURES/9

96. Bhattacharya S, Dunn P, Thomas CG, et al. ImmPort, toward repurposing of open access immunological assay data for translational and clinical research. Sci Data. 2018;5. doi:10.1038/SDATA.2018.15

97. Lin GL, Nagaraja S, Filbin MG, Suvà ML, Vogel H, Monje M. Non-inflammatory tumor microenvironment of diffuse intrinsic pontine glioma. Acta Neuropathol Commun. 2018;6(1):51. doi:10.1186/S40478-018-0553-X/FIGURES/4

